# Ligand-induced conformational changes in the β1-Adrenergic Receptor Revealed by Hydrogen-Deuterium Exchange Mass Spectrometry

**DOI:** 10.1101/2024.02.07.579309

**Authors:** Joanna Toporowska, Parth Kapoor, Maria Musgaard, Karolina Gherbi, Kathy Sengmany, Feng Qu, Mark Soave, Hsin-Yung Yen, Kjetil Hansen, Ali Jazayeri, Jonathan T.S. Hopper, Argyris Politis

## Abstract

G-Protein Coupled Receptors (GPCRs) constitute the largest family of signalling proteins responsible for translating extracellular stimuli into intracellular functions. When dysregulated, GPCRs drive numerous diseases and are the most targeted proteins in drug discovery. GPCR structural dynamics and activity can be modulated by a wide range of drugs, including full/partial agonists and antagonists. While crucial for developing novel therapeutics targeting GPCRs, the structural dynamics of the receptors associated with their activity upon drug interactions are not yet fully understood. Here, we employ Hydrogen Deuterium Exchange Mass Spectrometry (HDX-MS), to characterise the structural dynamics of turkey β1-adrenergic receptor (tβ1AR) in complex with nine ligands, including agonists, partial agonists and antagonists. We show that dynamic signatures across the GPCR structure can be grouped by compound modality. Surprisingly, we discovered repeated destabilisation of the intracellular loop 1 (ICL1) upon full agonist binding and stabilisation upon antagonist binding, suggesting that increased dynamics in this region are an essential component for G-protein recruitment. Multiple sequence alignments and molecular dynamics simulations indicate that L72 in ICL1 plays important structural role. Differential HDX-MS experiment of tβ1AR and tβ1AR L72A construct in complex with miniGs, in response to various ligands, suggests involvement of ICL1 in stabilising the GDP bound state by influencing the stability of HG helix of miniGs. Overall, our results provide a platform for determining drug modality and highlight how HDX-MS can be used to dissect receptor ligand interaction properties and GPCR mechanism.

**Significance statement:** Recent advances in hydrogen-deuterium exchange mass spectrometry have allowed probing conformational signatures of challenging membrane protein assemblies. We studied the structural dynamics of a class A GPCR, namely tβ1AR, in response to diverse ligands including agonists, antagonists and partial agonists. We demonstrate that the functional effect of compounds can be discerned by simply profiling the dynamics induced across the receptor, without the need for downstream interaction partners. We showed that ICL1 undergoes a significant change in dynamics between activated and inhibited states consistent with a role in downstream signaling pathways in class A GPCRs.

## Introduction

G-protein coupled receptors (GPCRs), a superfamily of more than 800 unique members, and the largest family of receptor proteins in mammals, are responsible for a diverse range of physiological responses^1,2^. They share a common structural arrangement via seven transmembrane (7TM) helices that are bound by three intracellular (ICL) and three extracellular (ECL) loops. GPCRs respond to various extracellular stimuli, including hormones, metabolites and neurotransmitters^1^. Ligand binding to GPCRs serves to induce, or stabilise, distinct conformational arrangements that define the intracellular physiological response. Canonical agonist binding, for example, results in GPCR conformations that selectively allow the receptor to recruit and activate specific heterotrimeric G-proteins, resulting in the generation of second messenger molecules. Additionally, some ligands induce a functionally selective (or “biased”) response, driving signalling through either specific G-proteins or through other transducers such as β-arrestins. These differences in signalling pathways, which essentially define the physiological action of a particular ligand, are driven by allosteric conformational changes of the receptor. This highlights the challenges in future GPCR drug discovery, where early knowledge of the ligand-induced conformational dynamics of the receptor can greatly improve the identification of early hits with the correct pharmacological response, as well as drive correct decisions during the SAR development of those leads.

The β1-adrenergic receptor belongs to the class A, or rhodopsin-like, subfamily of GPCRs along with, β2AR and β3AR, which share more than 50% of sequence similarity^5^. β1AR is abundantly expressed in cardiac tissue and is critical for the physiological regulation of heart rate. Antagonists of β1AR are often referred to as β-blockers and inhibit binding of endogenous ligands norepinephrine and adrenaline to the catecholamine pocket, ultimately preventing receptor activation and resulting in a reduction in heart rate^3,15^. Structurally most of the transmembrane region is highly conserved among all GPCRs. The remarkable differences in the sequence are mostly present in the ICL and ECL regions which allows the wide range of ligands to bind to the GPCR orthosteric site and regulate receptor activity^1^. Mechanisms and structural transitions behind receptor activation are still unclear.

Hydrogen Deuterium Exchange (HDX) has emerged as powerful tool for monitoring the exchange of backbone hydrogen atoms for heavier deuterium in bulk D_2_O over different time intervals^9,10,12^. The rate of exchange depends on the local environment around a given backbone hydrogen atom, as well as the hydrogen bonding and accessibility of the local segment in which the specific hydrogen atom is located. Thus, measuring the rate of exchange can reveal details about local structural changes and dynamics. Here, deuterium uptake across the protein is determined by proteolytically digesting the protein and subsequently using mass spectrometry (MS) to measure the mass changes present across the identified peptides^12^. Recent workflows developed in our group and others have enabled experimental measurements of integral membrane proteins which achieve high sequence coverage^13,14,15^ and allow a more comprehensive analysis of dynamic conformation changes in these targets. In particular, HDX-MS offers advantages over traditional approaches; it tolerates heterogenous environments (e.g., lipid nanodiscs), has low sample requirements and there is no need for chemical probes or bio-orthogonal labels^12,17^. Despite advances and early successes^18,19^, interrogating the conformational dynamics of GPCRs remain a challenging task, primarily owing to difficulties in solubilising functionally competent preparations of these important biomolecules. To overcome that issue and efficiently solubilise membrane proteins, many different solutions or methods have been proposed including surfactants (e.g., detergents) or nanodiscs (lipid-based environment) and SMALPs (styrene-maleic acid lipid particle technology)^20^. They provide a “protective layer” surrounding the transmembrane part of the protein. However, the resulting heterogeneous mixtures may complicate the experiments, leading to extra steps in the optimisation protocol (e.g., removal of lipids)^12^. Here we have successfully optimised the use of HDX-MS to be able to interrogate the structural dynamics of tβ1AR^32^.

Using differential HDX-MS, we have monitored the global confirmational dynamics of the receptor in response to a diverse range of ligands which induce defined pharmacological responses to receptors. Our findings illustrate distinct dynamic signatures across the receptor upon full and partial agonist, and antagonist binding. Interestingly, opposing dynamics were consistently found in the ICL1 loop, appearing to highlight a structurally essential component in driving receptor activation vs. deactivation. This prompted us to further explore the structure and function of ICL1 using site-directed mutagenesis in combination with molecular dynamics simulations and cell-based functional assays. We provide a peptide-level characterisation of this loop and reveal the impact of ICL1 on downstream signalling in a class A receptor.

## Results and Discussion

### HDX-MS optimisation for high peptide coverage

To date, the study of GPCRs by HDX-MS has been frustrated by low/medium sequence coverages. Therefore, in our first step, we optimised conditions to achieve high sequence coverage for tβ1AR. Initially we embarked on screening against various quench buffer compositions as well as different pepsin columns (**Supplementary Figure 1**). Optimal sequence coverage was achieved using a self - packed pepsin column with Immobilized Pepsin Agarose, which allowed us to achieve a sequence coverage of 75%. We observed that increasing DDM ((n-dodecyl-β-D-maltopyranoside) concentration in quench buffer from 0.01% to 0.1%, approximately 10x the documented CMC (critical micelle concentration), resulted in a significant increase (∼20%) in sequence coverage. It has been reported that β1AR contains disulphide bonds which can negatively impact the efficiency of enzyme digestion^16^. We maintained a TCEP (tris(2-carboxyethyl)phosphine hydrochloride) concentration of 100mM to reduce disulphides and give the optimum sequence coverage. Finally, we optimised the digestion to utilise a dual protease type XIII/pepsin column. The protease type XIII (from *Aspergillus saitoi)* is a non-specific enzyme, and its cleavage sites are complementary to pepsin^22^. The column change, led to a significant increase in sequence coverage up to 94%, and redundancy of 8.27 (**Supplementary** Fig.2). With the optimal quench composition and digestion efficiency established, we progressed to full HDX-MS experiments.

### HDX fingerprint of β1AR bound to antagonist

Initially we applied our optimised HDX-MS method to monitor the changes in conformational dynamics induced upon binding of three β1AR receptor antagonists, namely cyanopindolol, carvedilol and carazolol. Cyanopindolol and carazolol are high affinity antagonists for both β1- and β2-adrenergic receptors^24^. Carvedilol is a specific β1AR biased ligand that preferentially stimulates the β-arrestin signalling cascade, while displaying antagonistic properties towards G-protein recruitment^25^. Our data confirms that carvedilol show unique properties by having slightly different dynamics in comparison to other antagonists that have been tested.

We conduct HDX-MS experiments across four time points (15 sec, 2 min, 30 min and 120 min) and we use excess ligand concentrations to achieve > 98% binding occupancy (**Supplementary Figure 4**). We observed that various regions of the receptor are protected from HDX for all three ligand-receptor complexes. This behaviour is somewhat expected and is related to the increased stabilisation of its secondary structure and reduced receptor activity (**Figure 1A**). The protection observed in TM5 is induced by binding of the ligand in the catecholamine binding pocket located between extracellular parts of TM3, 4, 5, 6 and 7. Previous crystallographic studies of antagonists/inverse agonist structures illustrate that binding is localised to the TM7 and TM5 interface, involving key interactions with residues Ser211, Ser212 and Ser215^27^. Our data shows protection from deuterium incorporation for the peptides corresponding to the expected binding regions of TM5. We observed additional protection on the tip of TM4 and ECL2, which contains a short α-helical segment, that has been shown to be a component of the ligand binding site, forming the so called “lid” which enables access to the orthosteric site^1^. It is believed that ECL2 plays a key role in early stages of ligand recognition and could impact the specificity of the binding^5^. Protection on the tip of TM4 spans across all the time points with similar difference in deuterium uptake (∼0.6 Da), whereas on ECL2 the effect manifests only at early 15 sec and 2 min time points, which corresponds to a higher flexibility of this loop region that forms transient interactions in the presence of antagonists. Our data also demonstrate that binding of antagonists induces stabilising effects on ECL3 and the tips of TM6 and TM7, which increase with time. Computational studies have previously predicted that a secondary, transient binding site is located on ECL2, ECL3 and TM7, which may explain protection in these regions^28^. In addition to protection on the extracellular side of β1AR, we unexpectedly observed repeated stabilising effects on ICL1 for all tested antagonists, which to our knowledge has not been reported previously. Only carvedilol displays additional protection on the kink of TM2 that is located between TM1 and 7, which could be explained by its larger chemical structure containing the additional 3,4-dimethoxyphenethyl group, that extends beyond the catecholamine binding pocket in the direction of extracellular ends of TM1, 2 and 3 (**Figure 1B**). As mentioned above, residues located in this secondary binding cavity could be responsible for receptor selectivity of bigger ligands as well as to play a role in biased signalling^28^. Studies conducted on the angiotensin II receptor type 1 (AT1) receptor with a biased ligand demonstrate that its ability to signal through the β-arrestin pathway highly depends on the conserved proline located on the end of TM2^29^. Most significantly, our data demonstrate that all the antagonists investigated result in a reproducible global reduction in receptor dynamics. This suggests that compounds which have a common pharmacological modality can be identified in the absence of cell-based assays or downstream binding partners, simply by monitoring the changes in conformational dynamics induced by ligand binding. To test this hypothesis further, we expanded these studies to investigate receptor dynamics in the presence of agonists.

**Figure 1.**
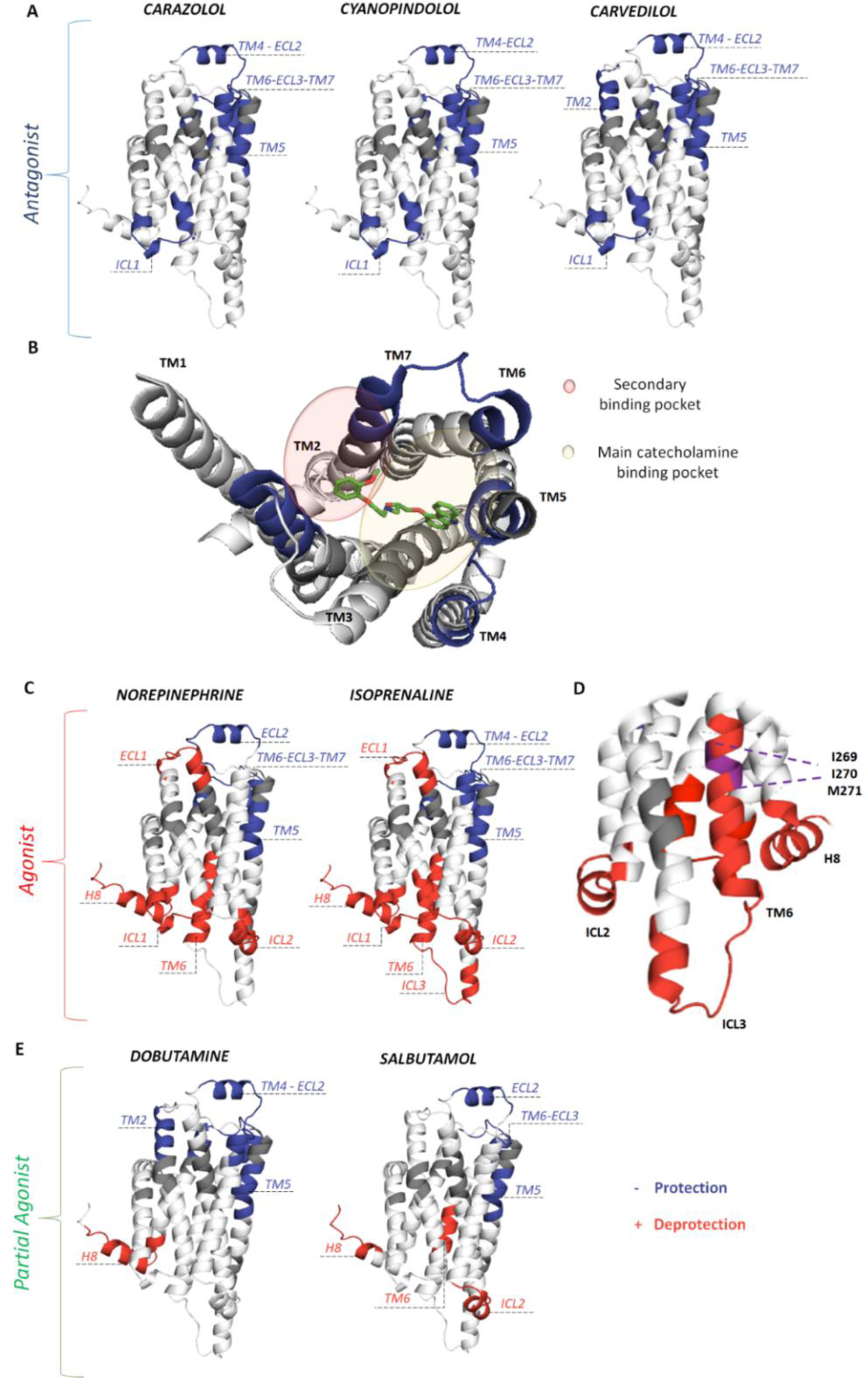
Comparison of HDX of tβ1AR apo state and tβ1AR bound to all tested ligands. Regions showing significant differences in HDX for tβ1AR bound to antagonist carazolol, cyanopindolol and carvedilol **A**, agonist norepinephrine and isoprenaline **C**, and partial agonist dobutamine and salbutamol **E** mapped onto the β1AR crystal structure (modelled by the use of PDB structures; 2VT4 chain A and 6IBL chain A). Blue colour indicates protection from the exchange and red deprotection. **D** Intracellular view of β1AR bound to isoprenaline with displayed deprotection onTM6 observed in our data, purple colour indicates residues (I269, I270, M271) that are most likely responsible for destabilisation of TM6. **B** Extracellular view of β1AR bound to carvedilol (PDB: 4AMJ) with displayed protection onTM2 observed in our data. The approximate areas of main catecholamine binding pocket and secondary binding pocket are highlighted in yellow and red respectively. We can see the additional 3,4-dimethoxyphenethyl group in the carvedilol structure extending beyond the main catecholamine binding pocket into the secondary binding site, explaining the observed protection on TM2.

### HDX fingerprint of β1AR bound to agonist

We next characterised the conformational effects of two agonists, isoprenaline and the endogenous ligand of β1AR, norepinephrine. Previous structural studies illustrate that agonists bind to the same β1AR catecholamine binding pocket as antagonists, however, there are additional rotamer conformational changes of Ser212 and Ser215 for agonist^23,28^. Similar to antagonists, agonist binding induced protection on TM5, TM4-ECL2 and TM6-ECL3-TM7, which sits directly within the orthosteric site and contains residues essential for binding. In contrast to antagonist binding, we observed common deprotection, indicating increased receptor dynamics, in multiple regions of the receptor (ICL1, ECL1, ICL2, ICL3, TM6 and H8) (**Figure 1C).** However, there are also some differences between the two agonists, including protection on the tip of TM4, and deprotection on ICL3 observed for isoprenaline but not for norepinephrine. It is well established that in all GPCRs intracellular loops play an important role in receptor activation, selectivity, and specificity^5^. Many studies report that coupling of GPCR to G-protein takes place through the cytoplasmic loops with ICL3 being of particular interest as it is responsible for determining specificity between different G-proteins^30^. GPCRs couple to G-protein at the cytoplasmic side mostly involving the end of TM3/ICL2, TM6 and TM5/ICL3^31^. Recent native MS studies state that isoprenaline induced β1AR complex formation with miniG_i_, whereas no complex formation was observed with norepinephrine^32^. Consistent with these findings, we have observed increased deuterium uptake on ICL3 for isoprenaline only, which highlights its ability to act as biased agonist, as well as the crucial role of ICL3 in defining specificity for distinct G-proteins and facilitating biased signalling pathways. In contrast to ICL3, enhanced dynamics of ICL2 are likely related to the strength of G-protein coupling rather than specificity^5^. It has been suggested that ICL2 acts as a switch allowing G-protein binding and highlighting its interaction with receptor conserved DRY motif located on TM3^5^. Increased deuterium uptake for both loops suggests that they play fundamental roles in recruitment of G-proteins and ultimately in receptor activation. Our data also reveals deprotection on TM6 and the cytoplasmic end of TM5 for both ligand-receptor complexes with similar level of deuterium incorporation (∼0.6Da for TM6). This effect supports a large outward movement of TM6 and TM5 upon agonist binding, confirming that those helices undergo significant conformational changes during receptor activation^33^. Notably, the effect on TM6 is only observed for the peptides containing three non-polar amino acid residues; two Isoleucine (I269, I270) and Methionine (M271) located closely to the core of TM6 (**Figure 1D).** This observation likely underlines the importance of those residues in so called “pivotal” movement of TM6 around its centre, where the intracellular side of the helix moves outwards, and the extracellular side moves inwards closer to the binding pocket^34^. Unexpectedly, we found that a unique characteristic of both agonists was their capability to induce increased dynamics on ICL1, opposite to antagonists which induced decreased dynamics on ICL1.

### HDX fingerprint of β1AR bound to partial agonist

Interesting candidates to expand the applications of our approach were dobutamine and salbutamol which are characterised as high affinity partial agonist for β1AR and β2AR, respectively. For both partial agonist-tβ1AR complexes we could observe similar conformational changes on the extracellular face of tβ1AR (ECL2, TM5, TM6-ECL3-TM7) as we observed for our agonist and antagonist bound structures (**Figure 1E**). On ECL2 we could see significantly bigger protection for antagonist than agonist or partial agonist (∼1.2 Da, 0.5Da, 0.4Da respectively). On TM5 we noted various deuterium uptake differences between the ligands, with all tested antagonists inducing protection of ∼2Da, isoprenaline ∼1.3Da, norepinephrine 0.5Da, dobutamine 1.9Da and salbutamol 0.7Da. It is interesting to speculate that this variety in deuterium uptake between the different ligands is likely linked to affinity of the compounds to the receptor (**Supplementary Figure 4**). Protection on TM6-ECL3-TM7 does not vary much between antagonist, agonist, and partial agonist with ∼0.6Da, 0.5Da, 0.4Da uptake difference respectively (averaged values for all ligands). For dobutamine, we also observed additional stabilisation induced on the tip of TM2. Dobutamine is a considerably larger ligand, and once bound to the receptor would extend beyond the catecholamine binding pocket, causing protection on TM2, similar to our data recorded for the biased agonist carvedilol. TM2 has also been recognised by NMR studies as an important location indicative of distinct receptor conformations, including carvedilol-bound state^35^. Similar to norepinephrine, salbutamol did not induce protection on the tip of TM4, whereas all other tested ligands did. Most likely, those observations highlight the different modes of ligand binding in the orthosteric binding pocket and their ability to stabilise the extracellular site of the receptor. Notably, the pharmacophore moiety of salbutamol docks in the catecholamine binding pocket in the same manner as endogenous agonist norepinephrine^36^, which possibly explains the lack of protection on the tip of TM4 for both ligand-receptor complexes. The major differences between those two partial agonist-receptor complexes were noted on the cytoplasmic site of tβ1AR. Salbutamol-bound tβ1AR is considerably more dynamic than dobutamine-bound tβ1AR (deprotection only on H8) inducing additional deprotection on ICL2 and TM6, however, still less than the agonist-receptor complex, which displays the most dynamic state of the receptor. Deprotection on TM6, with the average statistically significant difference in deuterium uptake (120 min time point) of ∼0.47Da, ∼0.59Da and ∼0.76Da for salbutamol, norepinephrine and isoprenaline respectively, indicates smaller outward movement of TM6 for salbutamol-receptor complex. The number of peptides displaying the effect on TM6 was also significantly lower for salbutamol. This is potentially indicative of an intermediate active state of the receptor related to salbutamol’s partial agonism^36^. Observed differences between dobutamine and salbutamol might be linked to variation in affinity between those two partial agonists (salbutamol: *k*_*d*_ = 20.89 *μM*, dobutamine: *k*_*d*_ = 5.88 *μM*)^31^, and they both could propagate slightly different dynamics across the receptor spanning from binding pocket to cytoplasmic site of the receptor. Crystallographic studies of the β1AR in complex with agonist and partial agonist reveal that the difference in binding between those two classes of drugs is a reduction in the number of hydrogen bonds between the ligand and the residues located in catecholamine binding pocket, whereas motifs in the intracellular site of the receptor were almost the same^23^. In addition, partial agonists do not alter the conformation of Ser215^23^. It is worth noting that there were no effects observed on ICL1 for both partial agonists. Our data shows clear differences between the receptor dynamics of antagonist-, full agonist-, and partial agonist-bound tβ1AR, information that is ultimately lost in structure determination through x-ray crystallography or cryo-electron microscopy. Such dynamics provide further insight in understanding receptor dynamics and responses to drugs with different pharmacologies, therefore illustrating the power of combining multiple complementary approaches for understanding function.

### The role of ICL1 in receptor activation

Our studies of the conformational changes of tβ1AR in response to various ligands, have repeatedly highlighted the increased dynamics of intracellular loop 1 (ICL1) in the presence of full agonists. To explore the role of ICL1 in receptor activation or G-protein recruitment we extended our HDX-MS experiments to include two additional ligands, derivatives of full agonist isoprenaline; colterol hydrochloride, which maintains a level of coupling activity similar to isoprenaline, and orciprenaline, which has significantly reduced ability to induce β1AR-G protein recruitment^32^. Our data reveals increased dynamics of ICL1 in the presence of colterol hydrochloride, similar to the effect observed for the full agonist isoprenaline, however, no significant effect was observed in the presence of orciprenaline (**Figure 2**). These results suggest that increased dynamics in ICL1 are only established upon full activation of the receptor and may play an essential role in allowing agonists to transduce signalling down the G-protein pathway.

**Figure 2.**
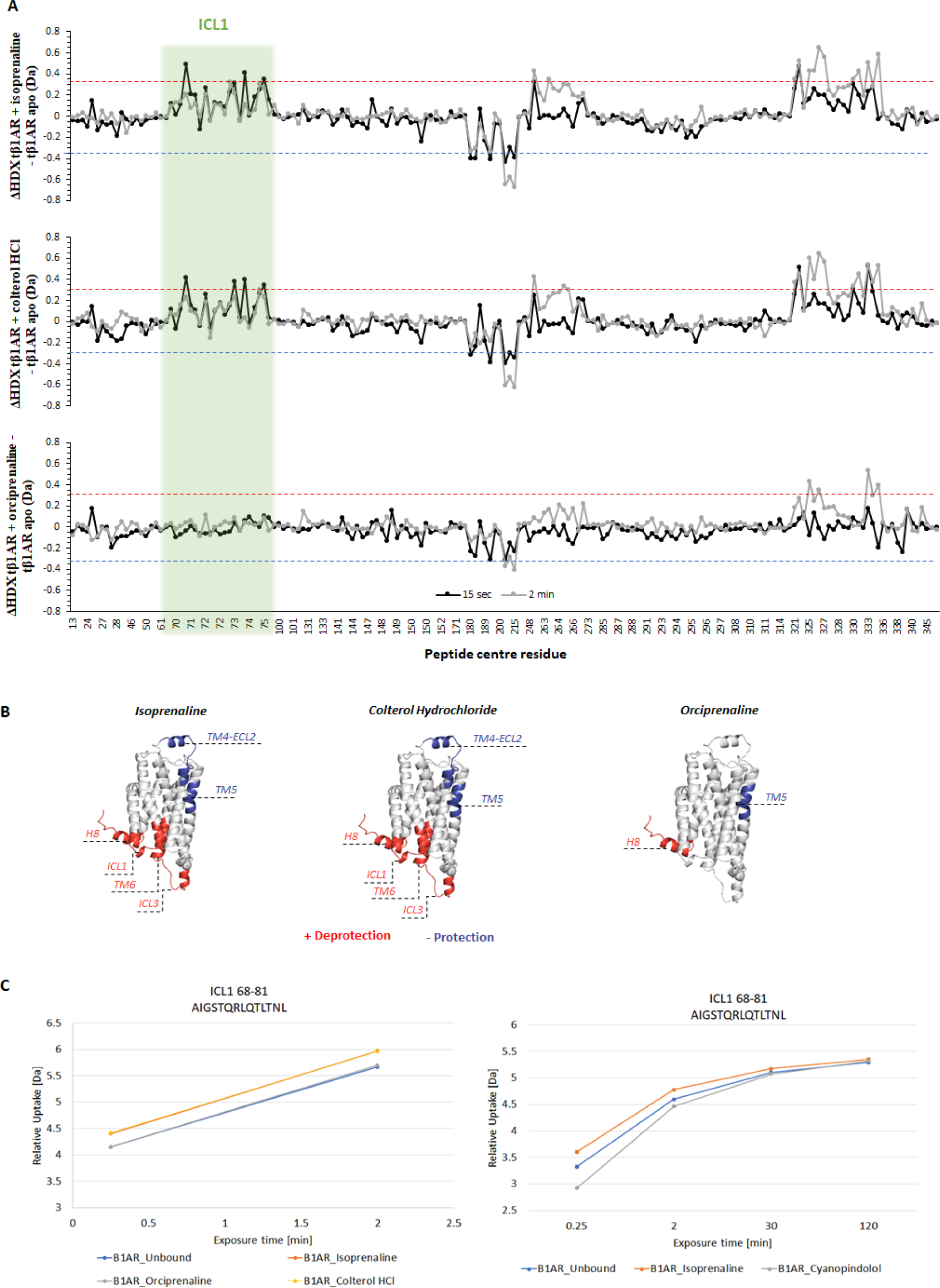
Comparative HDX analysis of tβ1AR apo state and tβ1AR bound to Isoprenaline, Colterol HCl and Orciprenaline. **A** Butterfly plots for HDX-MS experiment for tβ1AR with isoprenaline derivatives. These plots display differences in relative deuterium uptake (Da) between tβ1AR alone (apo) and tβ1AR+isoprenaline, tβ1AR+colterol HCl, and tβ1AR+orciprenaline, respectively. The black line indicates 15 sec time point and grey line 2 min time point. The blue and red dotted lines set at ∼0.3 Da indicate the threshold for significant differences in deuterium uptake calculated by Deuteros^33^. The ICL1 region is highlighted in green. It clear that all ICL1 containing peptides display the destabilisation effect for isoprenaline and colterol HCl, and no peptides showing any effect in orciprenaline bound state. **B** Regions showing significant difference in HDX for tβ1AR bound to isoprenaline and its derivatives, mapped onto β1AR crystal structure (modelled by use of PDB structures; 2VT4 chain A and 6IBL chain A). **C** Uptake plots displaying the effect on ICL1 for two separate experiments; isoprenaline derivatives and isoprenaline/cyanopindolol respectively.

To investigate the influence of ICL1 on receptor function, we performed mutational analysis in the ICL1 region of tβ1AR. The central part of ICL1 is composed of four residues in tβ1AR; Q70, R71, L72 and Q73. First, we generated a sequence logo (**Figure 3A**) based on multiple sequence alignments for the top 1000 hits from a Blast search on the wild-type turkey β1A sequence. The sequence logo illustrated that residues R71, L72 and Q73 (β1AR numbering) are well conserved. It is evident from the structure of tβ1AR that ICL1 is in close proximity of the horizontal H8 (**Figure 3B**). Inspecting potential direct interactions between ICL1 and H8 (**Figure 3C**) revealed that the lesser-conserved Q70 but also the more conserved Q73 point away from H8 and do not seem to form direct interactions with other structural elements of the protein. R71 of ICL1 could potentially form a salt bridge to D348 of H8, although their arrangement does not indicate a strong salt bridge. L72, on the other hand, seems to potentially pack into a small hydrophobic pocket formed especially by the C_α_ of D348 along with F349 and A352 of H8 (**Figure 3D**) but also with potential contributions from A65 in TM1 and F353 in H8. Especially F349 and F352 are well-conserved. We predicted that these interactions may be key for controlling the dynamics of ICL1. To disrupt this packing of ICL1 and H8 we mutated L72 to alanine. MD simulations were performed to assess the ICL1-H8 association in the apo state of both the tβ1AR turkey β1AR (modelled as PDB ID:4AMJ^50^) and a computationally constructed L72A mutant. Measuring the distance between ICL1 and H8 (as defined in **Figure 3E**) during simulations predict the ICL1-H8 association to be relatively stable in the tβ1AR apo state, but heavily disrupted in the L72A mutant, showing great fluctuations in the distance (**Figure 3F**).

**Figure 3.**
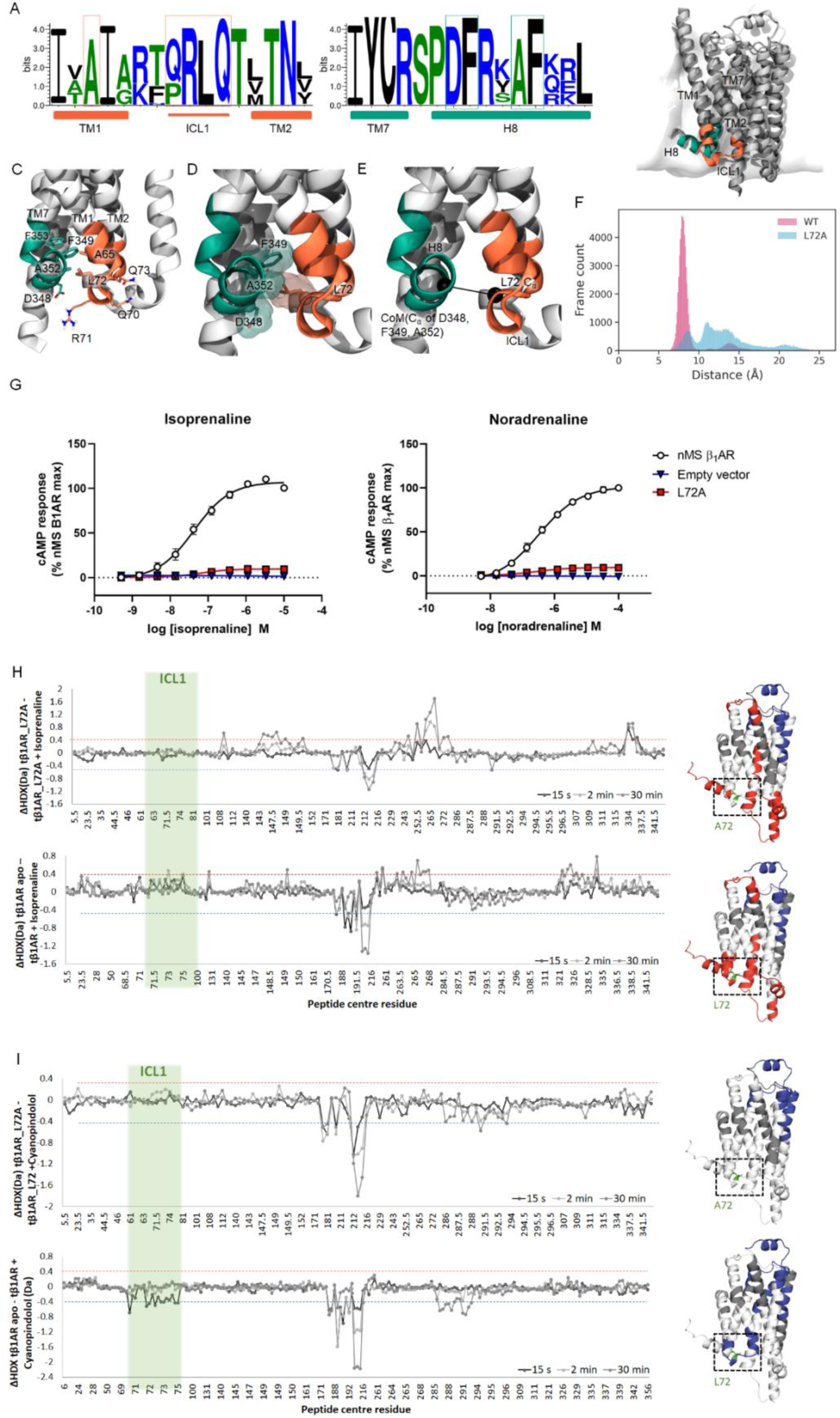
Design, dynamics and functional consequence of tβ1AR L72A. **A** Sequence logos for the 1000 top sequence from a blast search on the turkey WT β1AR sequence, showing the cytoplasmic end of TM1 and TM2 along with the connecting ICL1, as well as the cytoplasmic end of TM1 along with the H8 helix. Secondary structure elements are highlighted with the solid-coloured boxes below the plots. **B** Overall structure of β1AR (PDB: 4AMJ)^50^ illustrated in cartoon, placed in a POPC membrane model (white surface). ICL1 and H8 are highlighted using the same colours as the secondary structure bars in panel A. **C** An enlargement of the ICL1-H8 region with potentially important residues shown in licorice. The included residues are highlighted on panel A in thin boxes. **D** Same view as **C** but highlighting the packing of L72 against D348, F349 and A352. The C_α_ atoms are shown in vdW representation, and the side chains are shown with licorice as well as transparent surface. **E** Definition of the ICL1-H8 distance which is measured between the two black spheres defined by the C_α_ carbon atom of L72A and the centre of mass (CoM) of the C_α_ carbon atoms of D348, F349 and A352, respectively. **F** Distribution of ICL1-H8 distances as measured for a total of 1.2 μs of MD simulation for each of the tβ1AR (pink) and the L72A mutant (blue). **G** Isoprenaline and noradrenaline concentration response curves for cAMP accumulation determined in CHO cells transiently transfected with either empty vector, tβ1AR (nMS) or L72 tβ1AR. Data are expressed as mean ± SEM of 3-11 independent experiments performed in duplicate. Error bars not shown lie within the dimensions of the symbol. **H** Butterfly plots for tβ1AR L72A (top) and tβ1AR (bottom) bound to isoprenaline. **I** Butterfly plots for tβ1AR L72A (top) and tβ1AR (bottom) bound to cyanopindolol. Green colour highlights regions of ICL1. For tβ1AR L72A there is no deprotection on ICL1 upon isoprenaline binding as well as no protection upon cyanopindolol binding. Whereas both of those effects can be seen for tβ1AR upon respective ligand binding. Red and blue dashed lines indicate deprotection and protection limits respectively, calculated by Deuteros 2.0^33^.

We designed a mutated construct to allow us to examine the effects of the L72 residue in further experiments. We first determined the functional consequence of these mutations in CHO cells transiently transfected with tβ1AR and L72A tβ1AR by measuring cAMP production in response to isoprenaline and norepinephrine. In support of our predictions, the L72A variant showed a significantly impaired response to agonist, almost completely abolishing the resulting cAMP levels relative to parent tβ1AR for both tested agonists (**Figure 3G**). Receptor expression in membrane preparations of CHO cells transiently transfected with tβ1AR and L72A β1AR was determined using Western blot analysis, which confirmed that the L72A receptor was successfully expressed and not misfolded, but that lower expression levels were observed for the mutant compared to the parent receptor (**Supplementary Figure 6**). Taken together, these data suggest a potential key role of ICL1, and specifically the L72 residue, in propagating GPCR signaling, however, we cannot completely exclude that the drop in activity is due to a lower abundance of the mutant receptor. To further characterize the impact of the L72A mutation, the variant was purified and we conducted a differential HDX-MS experiment using the tβ1AR L72A mutant, in the presence of the agonist isoprenaline and the antagonist cyanopindolol. Interestingly, our data for the tβ1AR L72A mutant demonstrated no observable difference in deuterium uptake between the bound and unbound state of the mutant on ICL1, irrespective of whether an agonist or antagonist was present (**Figures 3H,I**). The interaction between ICL1 and H8 in class A GPCRs has been previously identified in a study investigating GPCR activation mechanisms. In the same research, the Leucine residue on ICL1 was recognized as a key player in establishing contact with H8^41^.

### Importance of β1AR intracellular loop 1 (ICL1) in allosteric activation of miniGs

Upon determining the influence of ICL1 on the conformational dynamics of tβ1AR, we then conducted differential HDX-MS experiments by coupling tβ1AR to miniGs and subsequently binding it to an agonist, antagonist, and partial agonist (**Supplementary Figure 15**). Our data suggest that all ligands induce a similar conformational pattern of the receptor when complexed with miniGs. Notably, we observed a protection on the extracellular side, which encompasses ECL1 and the apex of TM1 as well as ECL2 and TM5 - regions identified as ligand binding sites. The results also indicate a decrease in deuterium uptake on the cytoplasmic side of the receptor ICL2 and TM5/ICL3/TM6 in response to miniGs binding, corroborating previous structural studies on GPCR/miniGs complexes, thereby representing the primary contacts formed with miniGs^43^. We have also observed protection on ICL1 and H8, which we hypothesize contributes to the stabilization of the tβ1AR/miniGs complex by securing the receptor in a stable conformation (**Figure 4A**). The full agonist elicited larger differences in deuterium uptake in ICL2 and TM5/ICL3/TM6 in comparison to the partial agonist and antagonist. In addition, a significantly higher number of peptides displayed an effect in the isoprenaline-bound state, implying an enhanced stabilisation of the tβ1AR/miniGs complex upon agonist binding. In terms of miniGs, we noted a protection on H5, S2-S3, and H2 across all three complexes, which validates the central role of H5 as the interface establishing contact with β1AR. Intriguingly, Isoprenaline (agonist) stimulated increased deuterium uptake on H1, P-loop, and S5-HG, a phenomenon absent in the presence of the antagonist. The increased dynamics observed in the H1, and P-loop region can be attributed to a broken connection with H5 upon β1AR binding^46^. Furthermore, the observed deprotection on S5-HG and H1 is likely indicative of a disrupted contact between S5-HG, H1, and the nucleotide binding site, culminating in GDP release^46^. To investigate if the L72A mutation impact receptor/miniGs coupling, we performed differential HDX-MS experiment with the tβ1AR L72A mutant in the presence of miniGs and agonist-Isoprenaline (**Figure 4A**). Our results demonstrate that the L72A mutant could still bind to miniGs, inducing the same conformational changes on the intracellular and the extracellular side as tβ1AR with the exception for ICL1 and H8. Previously observed protection on ICL1 and H8 for the tβ1AR in complex with miniGs, has been abolished by the presence of the L72A mutation. Relative deuterium uptake for ICL1 is significantly higher for the tβ1AR L72A mutant than for tβ1AR, indicating increased dynamics of the loop in L72A mutant (**Figure 4B**).

**Figure 4.**
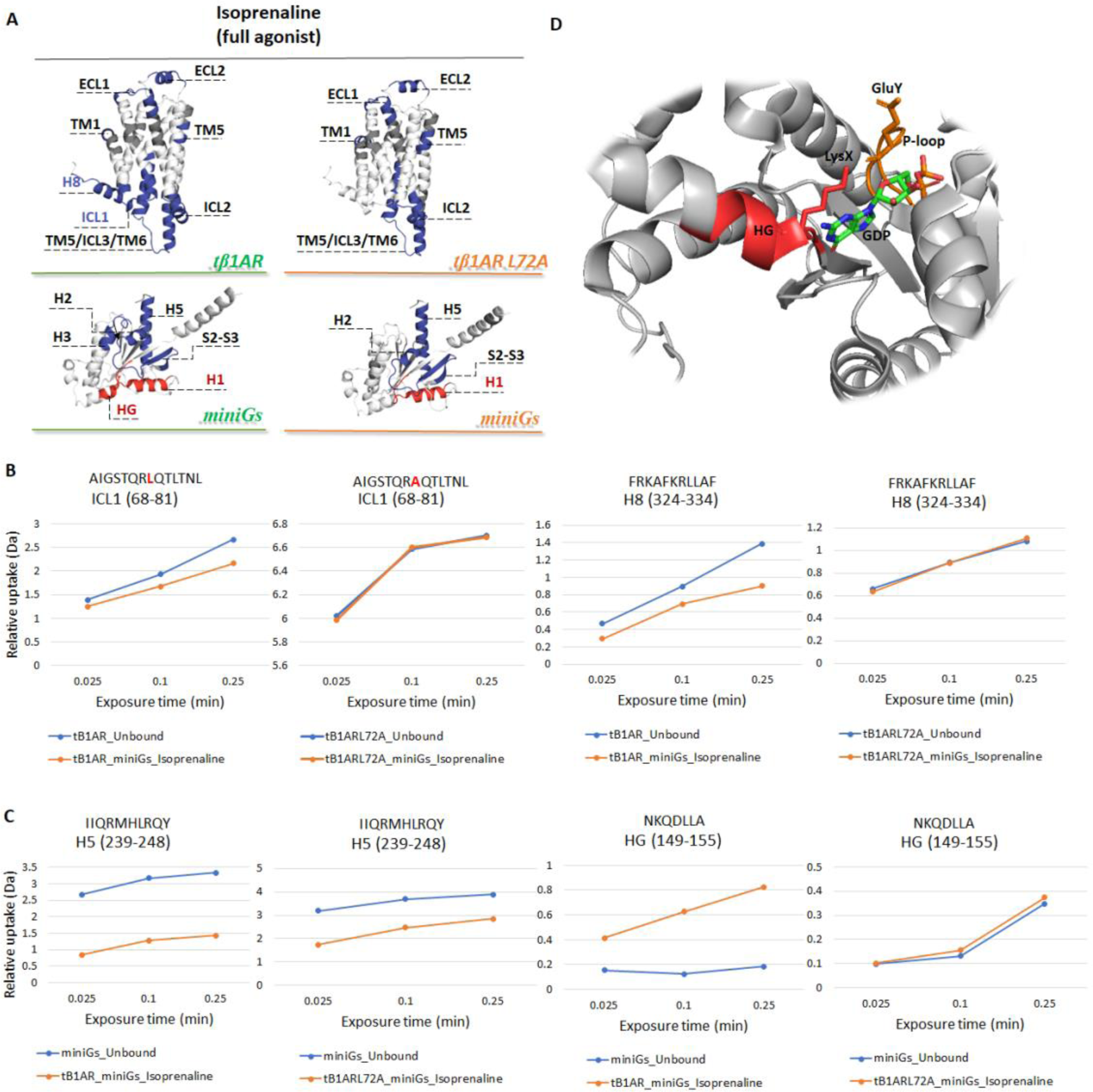
Results of differential HDX of tβ1AR/miniGs/various ligands complexes and tβ1ARL72A/miniGs/isoprenaline complex. **A** Regions showing significant difference in HDX for tβ1AR/miniGs complexes with isoprenaline (under green line) and tβ1AR L72A/miniGs complex with isoprenaline (under orange line) mapped onto β1AR crystal structure (modelled by the use of PDB structures; 2VT4 chain A and 6IBL chain A) and miniGs crystal structure. **B** Uptake plots displaying the effect on ICL1 and H8 for two independent experiments; tβ1AR/miniGs/isoprenaline and tβ1AR L72A/miniGs/isoprenaline. **C** Uptake plots displaying the effect on H5 and HG of miniGs for two independent experiments; tβ1AR/miniGs/isoprenaline and tβ1AR L72A/miniGs/isoprenaline. **D** The view of miniGs with bound GDP (PDB: 6EG8) and with highlighted residues (LysX, GluY) that are part of salt bridge between HG and P-loop.

Additionally, the mutation of L72A has abolished the increased dynamics of the S5-HG region of miniGs which was observed with the tβ1AR (**Figure 4C**). The level of protection observed on H5 of miniGs, which represents the primary binding epitope to the β1 adrenergic receptor (β1AR), is significantly reduced in the mutated L72A receptor. We hypothesize that the compromised protection on ICL1 and H8 in the tβ1AR L72A mutant may indicates dynamics that prevent the receptor from adopting a stable conformation that allows for all contacts with H5 of miniGs. As a result, there is a decreased difference in deuterium uptake on H5 observed in the coupling experiment with the L72A mutant. The interaction between H8 of class A GPCRs and H5 of miniGs, reported in the literature^47^could be affected in our experiments with the tβ1AR L72A mutant. Furthermore, a change in this interaction could affect contact re-organisation within miniGs, that are essential for its activation. Lack of deprotection on S5-HG could indicate that contact between H5 and S5-HG were not disrupted in the L72A complex as it is for tβ1AR. In the inactive state, residues located on HG of miniGs contact GDP, whereas in the active state (GPCR bound) this contact is disrupted to promote GDP release^46^. Studies performed on β2AR report that conformational changes on H5 of the Gα subunit affect the nucleotide binding region by directly impacting S5 and HG helix, which interact with guanine ring of GDP via two residues (LysX and AspZ)^48^. Increased disorder of those residues as well as increased mobility of HG were observed in GPCR – mediated activation. Additionally, one of these residues (LysX) located on HG is a part of important salt bridge formed between HG and P-loop (GluY) (**Figure 4D**), and whose disruption is a key element facilitating the GDP release^49^. Due to lack of deprotection on HG, our data demonstrate that contact between HG and GDP was not compromised in our experiment with tβ1AR L72A mutant indicating that the L72 residue on ICL1 of tβ1AR have an impact on stability of the GDP-bound state by directly affecting the dynamics of HG helix of miniGs

## Conclusions

Here, we have demonstrated the utility of HDX-MS approach in defining a conformational fingerprint of tβ1AR. We demonstrate that such fingerprints can be used to reveal the allosteric modulation of receptor dynamics in response to stimuli. We have exemplified this method by monitoring the change in dynamics induced by a range of compounds, such as full/partial agonists and antagonists. Our results highlight that compounds with similar pharmacological function impose, or stabilise, similar conformational signatures across the receptor, thereby allowing the mode of action of compounds to be discerned in a single experiment. This provides highly valuable insights for pharmaceutical research, allowing early hits which have the desired pharmacology to be progressed and allows more informed decisions as compounds undergo further SAR development towards candidate drugs. This is particularly important for drugs which are being developed to invoke a biased receptor response, whereby the development of any lead compound must maintain a strict propensity towards the desired signalling pathway. This gives further insights into structural aspects which are still widely debated, such as partial agonism. To exemplify this, biased ligands such as isoprenaline displayed reproducible deprotection on ICL3 whereas norepinephrine did not, suggesting that ICL3 deprotection could be used as a marker to confirm biased activity.

The findings of this research underscore the crucial role of Intracellular Loop 1 (ICL1) in receptor activation. Previous mutagenic analysis of residues in ICL1 of class B1 GPCRs has demonstrated that mutations in this loop region can significantly modulate receptor activation^38^. Concurrent molecular dynamic simulations of the Calcitonin Receptor-Like Receptor (CLR) revealed minor movement of ICL1, succeeded by rearrangements of ICL4 (the junction between TM7 and H8). Certain residues within ICL1 orientate towards H8, while residues of H8 are centrally directed into ICL1, suggesting a potential interaction between ICL1 and H8. These alterations are viewed as initial steps of receptor activation, occurring prior to movement of TM5 and TM6^38^. Further research on the active and inactive state structures of GPCRs from each class has reported that state–specific contacts are established not only to transmembrane helices but also to H8 and ICL1. Consequently, the active and inactive states of class A GPCRs are stabilized by rerouting contacts between residues located on transmembrane helices 1-7 as well as ICL1 and H8, with one of the residues identified as a state switch^41^.

In our studies of tβ1AR dynamics, we observed repeated destabilisation of ICL1 following binding of agonists. MD simulations, in combination with characterising residue conservation across several homologues for this loop, identified residue L72 as forming key interactions with conserved hydrophobic residues of H8. Mutational analysis in cell-based assays further support a potential functional importance of ICL1, essentially causing an almost complete loss of signalling in the single point variant. We cannot rule out that lower expression levels of the L72A may contribute to the magnitude of this observation. Nonetheless, HDX analysis using the L72A receptor variant revealed that conformational differences previously observed for tβ1AR ICL1 between agonist bound/unbound receptor were completely abolished. This suggests that mutation of this residue causes increased dynamics of the loop that results in contact disruption between ICL1 and H8. Studying the addition of miniGs into HDX experiments, clear protection at expected residues on the receptor confirmed that this variant is still able to form interactions with the G-protein. The protection previously observed for the tβ1AR on intracellular loop 1 (ICL1) and helix 8 (H8) has been compromised in the L72A mutant of tβ1AR. This suggests that the increased dynamics of ICL1 caused by the mutation disrupt the formation of contacts between the receptor and helix 5 (H5) of miniGs. As a result, the internal reorganization of contacts within miniGs, particularly between H5 and other areas of miniGs, may be significantly disrupted. Supporting this theory is the experimental observation that the tβ1AR L72A mutant does not show deprotection on the HG of miniGs that directly interacts with nucleotide binding site. We believe our data provides evidence that mutation of residue L72 on ICL1 of tβ1AR impacts GDP release and ultimately miniGs allosteric activation. Overall, our data provide insights into GPCR–G protein activation as well as GDP release mechanism.

## Materials and Methods

### Expression and purification of tβ1AR and tβ1AR_L72A mutant

Meleagris gallopavo tβ1AR construct (β114-E130W) harboured various thermostabilised point mutations and truncations at the N terminus, inner loop 3 and C terminus (β44-m23) previously described^32^ and an additional set of mutations to aid the purification step and retain functionality. The E130W mutation described here facilitated purification of the receptor in its apo state by stabilizing the intrahelical interactions between transmembrane helices TM3, TM4 and TM5. The constructs (tβ1AR and L72A mutant isoform) were over expressed in insect cells (sf9, Invitrogen, 11496015) using the Bac-to-Bac baculovirus expression system (Thermo Fisher). Expression vector pfastBac1 (Thermo fisher) was used to generate tβ1AR and mutant isoform (L72A) tβ1AR at a viral titre of multiplicity of infection of 0.5.

Briefly, cell membranes were prepared by lysing the cells in a cell disruptor (microfluidics; Quadro engineering) in 20mM Tris-HCl (pH8), 1mM EDTA, protease inhibitor (Roche) followed with subsequent wash steps using the ultracentrifuge. Membranes were finally resuspended in 20mM Tris-HCl (pH8) at a concentration of 10mg/ml.

The cell membranes were solubilized in 20mM Tris-HCl (pH8), 350mM NaCl, 3mM imidazole and 1.5% w/v n-dodecyl-β-d-maltopyranoside (DDM; Anatrace). The supernatant was obtained post the ultracentrifugation at 175,000 g for 1h and loaded onto a HiTrap TALON crude column (GE Healthcare) for affinity enrichment, pre-equilibrated in wash buffer 20mM Tris-HCl (pH8), 350mM NaCl, 3mM imidazole and 0.05% DDM. The receptor was eluted in presence of a gradient of 20mM Tris-HCl (pH8), 350mM NaCl, 250mM imidazole and 0.05% DDM. The protein was quantified on a SDS PAGE. The fractions containing receptor were pooled and concentrated to a final concentration of 2– 3mg/ml using an Amicon centrifugal filter with a molecular weight cut-off of 50 kDa for subsequent application.

### Expression and purification of miniGs

The engineered minimal G-protein, miniGs construct E392A was expressed in *E.coli* cells. The pellet from *E.coli* culture was resuspended in buffer A (40mM HEPES, pH 7.5, 100mM NaCl, 10mM imidazole, 10% v/v glycerol, 5mM MgCl_2_, 50μM GDP) with protease inhibitors, DNase I, lysozyme and DTT. The cells were lysed in the microfluidizer followed by lysate centrifugation at 38,000 x g for 45 min. Collected supernatant was filtered and loaded onto TALON column. The column was washed with buffer B (20mM HEPES, pH 7.5, 500mM NaCl, 40mM imidazole, 10% v/v glycerol, 1mM MgCl_2_, 50μM GDP). The protein was eluted with buffer B. Fractions containing the protein were concentrated and subjected to desalt with HiTrap desalting column in buffer D (20mM HEPES, pH 7.5, 100mM NaCl, 10% v/v glycerol, 1mM MgCl_2_, 10μM GDP). After desalting, protein concentration was measured and DTT, GDP and TEV protease were added to the sample in order to cleave of the histidine tag. Sample was incubated on ice for 2h, which was followed by reverse IMAC purification with TALON (cobalt) column equilibrated with buffer D. Protein was concentrated to the final concentration of 0.4mg/ml.

### Ligand solution preparation

Ligands were purchased from Combi blocks (carazolol), Enzo life science (cyanopindolol), Sigma-Aldrich (salbutamol and dobutamine), BLD Pharmatech (isoprenaline and orciprenaline) and Carbosynth (norepinephrine). Isoprenaline and orciprenaline were dissolved in water, whereas rest of the ligands were dissolved in 100% DMSO until the desired concentration was achieved. Concentration of DMSO was kept below 10% for all ligand-protein mixtures.

### Deuterium labelling of β1AR/ Ligand complexes

Purified β1AR at the concentration of 11μM was used for the experiment. The equilibration buffer (E) was composed of 20mM Tris – HCl, pH=8, 0.35M NaCl, 3mM imidazole, 0.05% DDM. The quench buffer (Q) was composed of 50mM K2HPO4, 50mM KH2PO4, 0.1% DDM, 100mM TCEP. The labelling buffer (L) was the same composition as (E) buffer except H_2_O was replaced with D_2_O (99.8%). The protein/ligand ratio was calculated based on (**Equation 1**) in order to enable around 99% binding occupancy of the receptor (**Supplementary Figure 4**)^44^. Deuterium labelling was performed by diluting 3.5 μl of protein (with or without ligand) in 32 μL labelling buffer. The protein sample was incubated for various time points (15sec, 2min, 30min and 120 min at room temperature) and then quenched with 35 μL buffer Q. Sample preparation (labelling and quenching) and injection was performed by an online robotic autosampler (LEAP HDX automation manager).

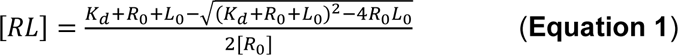

Parameters in the equation are as follow: *K*_*d*_ – dissociation constant, [*R*_o_] – concentration of the receptor before labelling, [*L*_o_] – concentration of the ligand before labelling, [*RL*] – fractional occupancy.

### Deuterium labelling of β1AR/miniGs/Ligand complexes

The coupling experiment was performed with β1AR and miniGs molar ratio of 1: 1.2 (11.5 μM: 13.8 μM). Sequence coverage obtained for miniGs was 92.4 % with redundancy: 7.26. The equilibration buffer (E) was composed of 20mM HEPES, pH 7.5, 100mM NaCl, 10% v/v glycerol, 1mM MgCl_2_, 10μM GDP. The quench buffer (Q) was composed of 50mM K2HPO4, 50mM KH2PO4, 0.1% DDM, 100mM TCEP. The labelling buffer (L) was the same composition as (E) buffer except H_2_O was replaced with D_2_O (99.8%) and 0.05% DDM was added. The protein/ligand ratio was calculated based on (**Equation 1**) in order to enable around 99% binding occupancy of the receptor (**Supplementary Figure 4**)^44^ Deuterium labelling was performed by diluting 3.5μl of protein (with or without ligand) in 32μL labelling buffer. The protein sample was incubated for various time points (15 sec on ice, 60 sec on ice and 15 sec in room temperature) and then quenched with 35μL buffer Q. Deuterium labelling and quenching of samples was performed manually on ice which was followed by manual injections.

### LC-MS workflow parameters for all HDX samples

Injected onto a nanoACQUITY UPLC system samples were digested with dual protease type XIII/pepsin column from NovaBioassays. HPLC run time was 11min at flow rate of 40μl/min with gradient. The columns used during the experiment were C18 trap (ACQUITY UPLC®BEH 1.7μm, Waters), a C18 column (ACQUITY UPLC®BEH, 1,7μm, 1.0 x 100 mm, Waters). For HPLC analysis, buffer A was composed of H2O with 0.1% formic acid, buffer B was acetonitrile with 0.1% formic acid at flow rate of 40μl/min. The gradient started with 8% B increased to 55% B within 8 min and then quickly increased to 85% B in 0.5 min, held for 1.5 min and decreased back to 8%. Analysis was performed by applying normal collision energy (CE) and look up table (LUT) collision energy, which is a time – dependent CE gradient. In this method the optimal collision energy is calculated for a set of peptides based on reference samples which has been shown to provide increased sequence coverage and redundancy for HDX experiments^21^. The mass range for MS was m/z 100-2000 in positive ion mode on the Synapt G2-Si mass spectrometer with ESI source and ion mobility cell, coupled to ACQUITY UPLC with HDX Automation technology (Waters Corporation, Manchester, UK). Leucine Enkephalin was applied for mass accuracy correction and sodium iodide was used as calibrant for the mass spectrometer. The trap and analytical column were kept at 0°C inside HDX manager. Between every injection clean blank was injected in order to remove any carryover. The data for each time point were obtained in three replicates. Data processing and analysis was performed processed using MassLynx (Waters), PLGS (ProteinLynx Global Server) used to analyse the MS data of unlabelled peptide and generate peptide libraries for each target protein. DynamX (Waters) used to analyse and quantify the deuteration for each peptide and Deuteros used to sort out statistically significant differences in deuterium uptake for peptides in two different conditions^32^.

### Multiple sequence alignment and sequence logos

The β1AR wild turkey sequence available on uniprot^51^ was used as reference sequence (uniprot ID P07700). A Blast search was performed through the uniprot website and the top 1000 hits were used as input for a multiple sequence alignment using Clustal Omega through EBI^52^. The resulting multiple sequence alignment was used for creating the sequence logos through WebLogo3^53^.

### MD simulations

Chain A from the 4AMJ^50^ pdb file of the turkey β1AR was used as a model for the tβ1AR apo state (resolution 2.30 Å). Both protein and water ascribed to chain B was deleted along with all other small molecules bound, apart from the sodium ion and the water molecules bound to chain A. The protein was prepared for simulation using the standard protein preparation workflow in Maestro^54^. The prepared protein structure was aligned to the 2y02 structure from the Orientations of Proteins in Membranes database^55^ to get a reasonable initial membrane position. An 80 Å x 80 Å pure POPC (1-palmitoyl-2-oleoyl-glycero-3-phosphocholine) bilayer was constructed around the protein using the Desmond^56,57^ System Builder and the system was solvated with SPC water molecules, neutralised and further ions were added to a concentration of 150 mM NaCl. The OPLS4 force field^58^ available in Maestro/Schrodinger was applied.

The single-point L72A mutation was introduced using the “Mutate” function in Maestro and the simulation system was constructed as described above for the tβ1AR system.

The systems were equilibrated in six steps, slowly increasing the temperature and reducing the force constants on the different components, generally following the default relaxation protocol for membrane protein systems in Desmond. Production runs likewise followed the default setup for Desmond simulations, with a temperature of 310 K and standard atmospheric pressure in the NPγT ensemble (constant particle number (N), pressure (P), lateral surface tension (γ), and temperature (T)). For each system 1 x 200 ns and 2 x 500 ns production runs were performed, starting from different seeds.

Analysis was performed using in-house Tcl scripts through VMD^59^ and structural images in panels 3B-3E were created using VMD. The plot in panel 3F was created using seaborn.

### cAMP accumulation assay

Chinese hamster ovary (CHO) cells from Merck (85051005) were maintained in DMEM/F12 cell culture medium supplemented with 10% fetal bovine serum and 1% L-glutamine, at 5% CO2, 37oC. Cells were grown to 70-80% confluency before transfection with β1AR constructs using FuGENE HD reagent (Promega) according to manufacturer’s instructions. The following day, transfected CHO cells were suspended in assay buffer (Hank’s balanced salt solution (HBSS) containing 5mM HEPES, 0.1% w/v BSA and 500mM 3-isobutyl-1-methylxanthine, pH 7.4) before incubation with increasing concentrations of noradrenaline or isoproprenaline for 1h at room temperature. After 1h, cAMP levels were measured using the cAMP Gs HiRange HTRF kit (Perkin Elmer) according to manufacturer’s instructions. Fluorescence resonance transfer (FRET) was detected on a PHERAstar plate reader (BMG) and FRET ratios used to determine cAMP levels from a cAMP standard curve from the same experiment. EC50 and Emax were estimated from concentration-response curves using a variable four-parameter logistic equation in Prism (v8; GraphPad).

## Acknowledgments

A.P. was supported by an EPSRC Fellowship (EP/V011715/1). This work was supported by EPSRC CASE Studentship co-funded by OMass Therapeutics to AP and JT.

## Author Contributions

J.T. and P.K. purified β1AR. J.T. purified miniGs J.T and A.P. designed the HDX-MS experiments, with input from J.T.S.H and H-Y.Y. J.T. carried out all HDX-MS experiments and data analysis. M.M. performed multiple sequence alignment, sequence logos and MD simulations. K.G. and K.S. performed cAMP accumulation assay. F.Q. and M.S. performed characterization experiments on transfected cells. K.H. helped in optimizing HDX-MS method. A.J., J.T.S.H. and A.P. supervised the project. J.T., J.T.S.H. and A.P wrote the paper with contributions from all authors.

