## Supplementary Information for "Ligand-induced conformational changes in the β1-Adrenergic Receptor Revealed by Hydrogen-Deuterium Exchange Mass Spectrometry"

### Supporting Figures

A

| Quench composition | Digestion coverage | Number of peptides | Redundancy |
|---|---|---|---|
| Waters pepsin + BPR+ 0.02% DDM | 30.4% | 18 | 1.60 |
| Waters pepsin + 0.02% DDM | 30.1% | 17 | 1.47 |
| Waters pepsin + 0.02% DDM, SD | 16.6% | 7 | 1.17 |
| Waters pepsin + 0.1%DDM | 50% | 29 | 1.62 |
| Waters pepsin + 0.02% DDM + 6M Gnd-HCl | 42.8% | 22 | 1.48 |
| Waters pepsin + 0.02% DDM + 6M urea | 43.1% | 23 | 1.63 |
| Waters pepsin + 0.1% DDM (8 to 55% LC) | 44.5% | 31 | 1.93 |
| Waters pepsin + 0.1% DDM + 400mM TCEP | 42% | 27 | 1.78 |

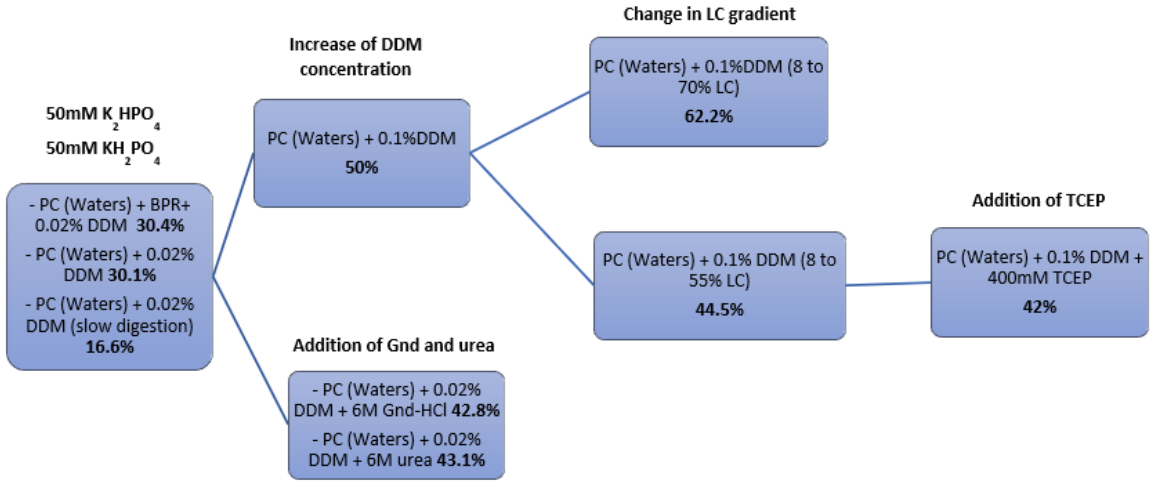


B

| Quench composition | Digestion coverage | Number of peptides | Redundancy |
|---|---|---|---|
| DIY PC + 0.1%DDM | 75.4% | 93 | 3.68 |
| DIY PC + CELUT + 0.1%DDM | 86.2% | 108 | 3.78 |
| DIY PC + 0.1%DDM, CE | 85% | 111 | 4.01 |
| DIY PC + 0.1%DDM, CE, SD | 85.5% | 112 | 4.06 |
| DIY PC+ 0.1%DDM, 15mM TCEP, CE | 74% | 72 | 3.18 |
| DIY PC + 0.1%DDM, 50mM TCEP, CELUT | 90.1% | 125 | 4.27 |
| DIY PC + 0.1% DDM, 100mM TCEP, CELUT | 90.1% | 142 | 4.97 |
| DIY PC+ 0.1%DDM_400mM TCEP, CELUT | 61% | 50 | 2.43* |
| DIY + 0.1%DDM, Rhizopus | 93.4% | 130 | 4.80 |
| DIY PC+0.1%DDM, 100mM TCEP, C8 | 85.1% | 120 | 4.31 |
| DIY PC + 0.1%DDM, 4M Gnd-HCl | 32% | 18 | 1.48 |
| DIY PC + 0.1%DDM, 6M Urea | 66% | 55 | 2.50 |
| Dual Protease + 0.1% DDM, 100mM TCEP, CELUT | 94% | 263 | 8.37 |

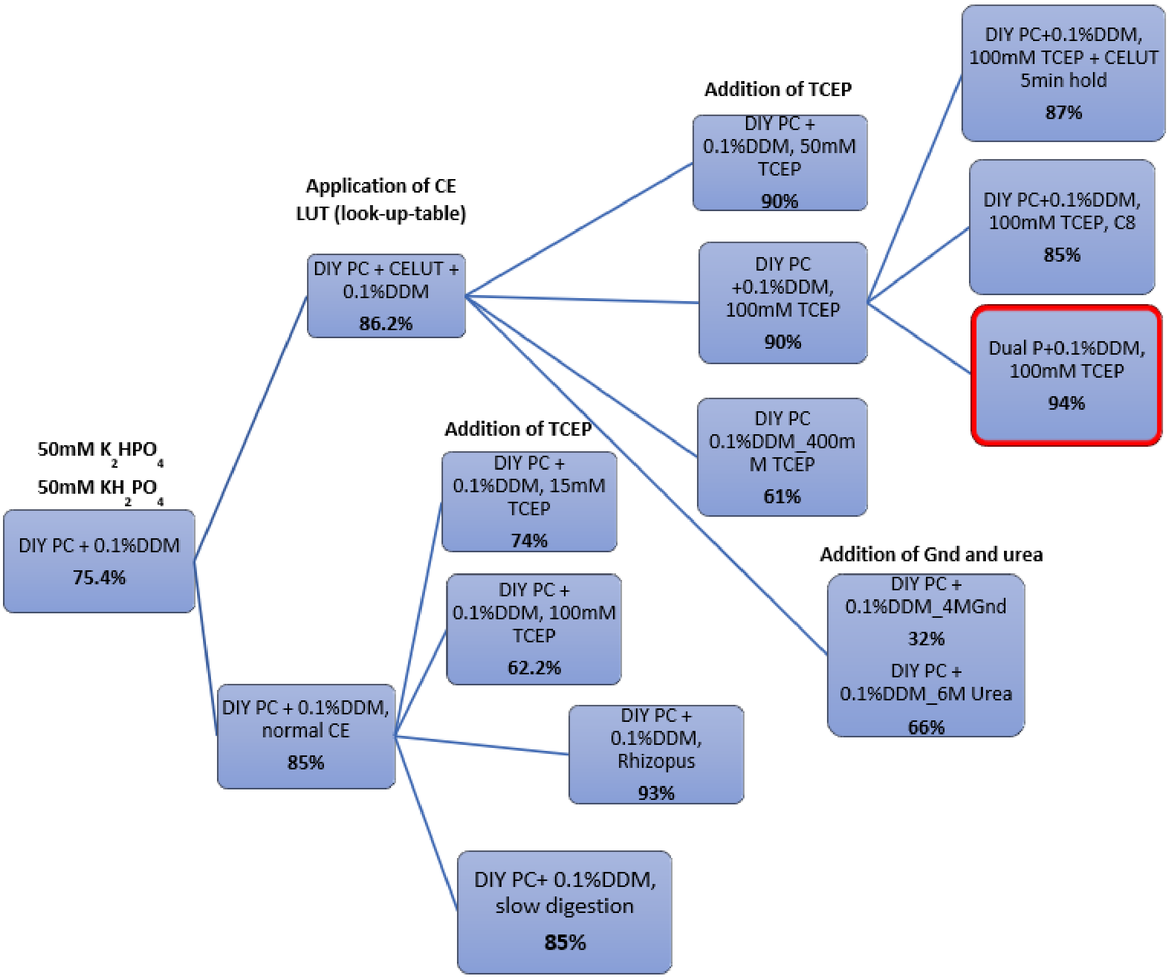


**Supplementary Figure 1.** Results, as measured by digestion coverage, number of peptides identified, and redundancy obtained from DynamX for tβ1AR quench buffer optimisation using the Waters Enzymate$^{TH}$ BEH pepsin column (**A**), self-packed pepsin column and dual protease type XIII (**B**). Red circle indicates final conditions, that have been applied to every HDX experiment. Abbreviations: BPR – back pressure regulator, SD – slow digestion, Gnd-HCL – guanidine hydrochloride, PC – pepsin column, DIY PC – self-packed pepsin column, CE – collision energy, CELUT – look up table collision energy, C8 – analytical column C8.

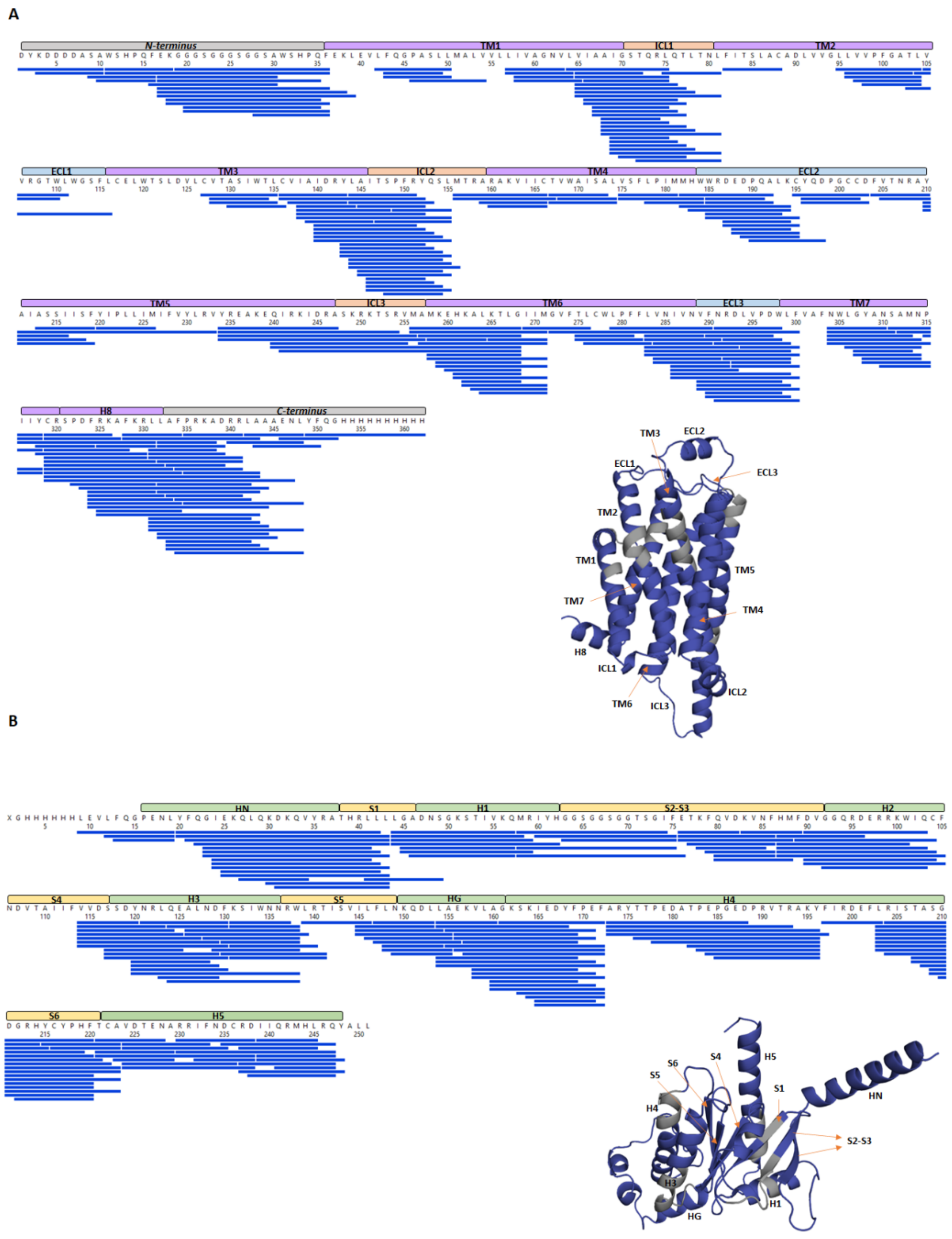

**Supplementary Figure 2.** Peptide maps obtained during optimisation step. **A** tβ1AR sequence coverage with displayed number of peptides covering parts of the protein. Achieved coverage: 93.6%, redundancy: 8.27. **B** miniGs sequence coverage 92.4% and redundancy: 7.26. Results are visualised in PyMOL. Blue colour indicates covered areas of the proteins.

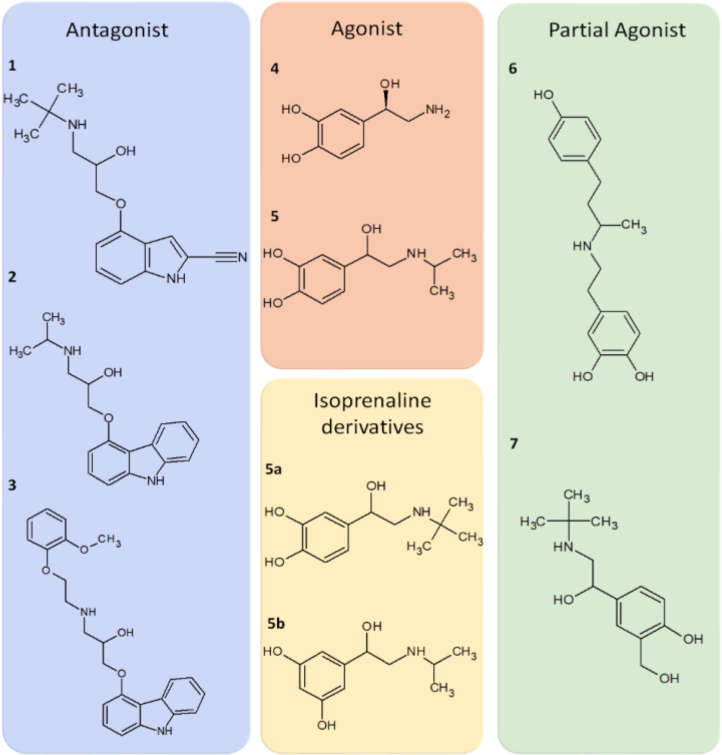


**Supplementary Figure 3.** Chemical structures of ligands used for analysis. Antagonists: **1** Cyanopindolol, **2** Carazolol, **3** Carvedilol. Agonists: **4** Norepinephrine, **5** Isoprenaline. Partial Agonists: **6** Dobutamine, **7** Salbutamol. Isoprenaline derivatives: **5a** Colterol Hydrochloride, **5b** Orciprenaline.

| Ligands | Kd [uM] | Ligand Concentration [uM] | Protein Concentration [uM] | % P bound |
|---|---|---|---|---|
| Isoprenaline | 0.870 | 300 | 11 | 99.70% |
| Norepinephrine | 1.819 | 300 | 11 | 99.37% |
| Dobutamine | 5.88 | 1000 | 11 | 99.41% |
| Salbutamol | 20.89 | 1820 | 11 | 98.86% |
| Cyanopindolol | 0.0000407 | 300 | 11 | 100% |
| Carvedilol | 0.004 | 300 | 11 | 100% |
| Carazolol | 0.000204 | 300 | 11 | 100% |

**Supplementary Figure 4.** Table displaying the properties of the ligands and protein used for experiments. The summary of the ligands Kd calculated based on the logKd values published by Jillian G Baker[31]. Ligand concentration was calculated in order to maximise the % of the ligand bound to the receptor in the labelling experiment. It consists of basic Kd calculations including the dilution factor, which in this case was equal to 9. Almost all ligands have high affinity to the receptor, apart from dobutamine and salbutamol. Therefore, concentrations of those ligands were significantly higher than others. Only salbutamol has slightly below 99% receptor occupancy due to its low solubility in DMSO and high Kd.

| | Antagonist | | | Agonist | | Partial agonist | |
|---|---|---|---|---|---|---|---|
| Motifs | Cyanopindolol | Carazolol | Carvedilol | Isoprenaline | Norepinephrine | Dobutamine | Salbutamol |
| ECL1 | | | | + | + | | |
| TM2 | | | − | | | − | |
| TM5 | − | − | − | − | − | − | − |
| TM6-ECL3-TM7 | − | − | − | − | − | − | − |
| ECL2 | − | − | − | − | − | − | − |
| TM4 | − | − | − | − | | − | |
| ICL1 | − | − | − | + | + | | |
| ICL2 | | | | + | + | | + |
| ICL3 | | | | + | | | |
| TM6 | | | | + | + | | + |
| H8 | | | | + | + | + | + |

**Supplementary Figure 5.** Summary table of all observed effects for tβ1AR in complex with various ligands. Table represents tβ1AR motifs where significant differences in deuterium uptake were observed, between tβ1AR – apo state and tβ1AR – ligand state. Blue coloured columns indicate antagonists, orange – agonists and green – partial agonists. Minus indicates protection from exchange and plus indicates deprotection.

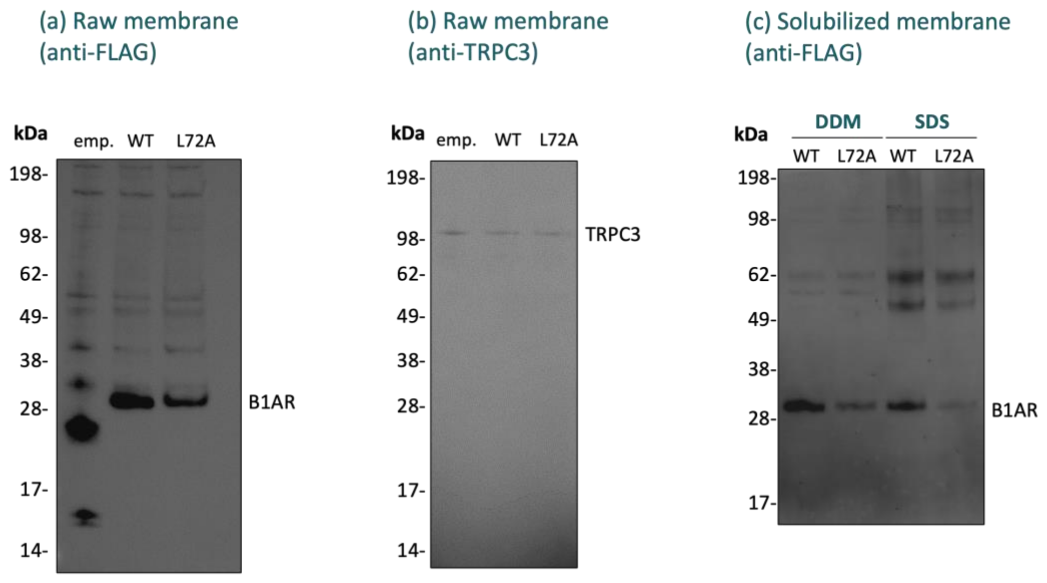


**Supplementary Figure 6.** Western blot analysis of membrane preparations from CHO cells transiently transfected with either an empty vector, a vector containing wildtype tβ1AR or L72A tβ1AR. (a) Expression was confirmed for both the wildtype and variant tβ1AR relative to transfection with an empty vector. Loading quantity was corrected using the BCA assay. (b) Western blot against the plasma membrane marker TRPC3 was also used as a loading control. (c) Crude membrane preparations of wildtype and variant tβ1AR were solubilised in either DDM or SDS, to confirm that the expression yields observed in (a) are due to folded protein (with only folded protein able to be solubilised by DDM[59].

#### tβ1AR apo vs tβ1AR + Carazolol

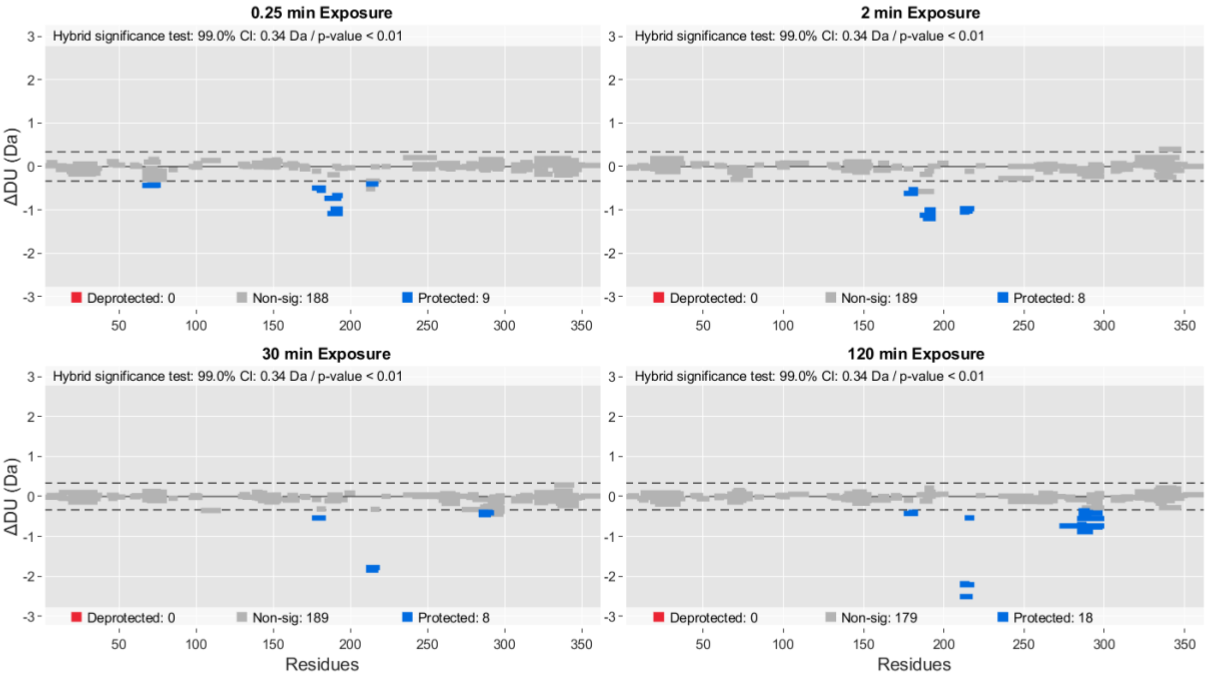


#### tβ1AR apo vs tβ1AR + Carvedilol

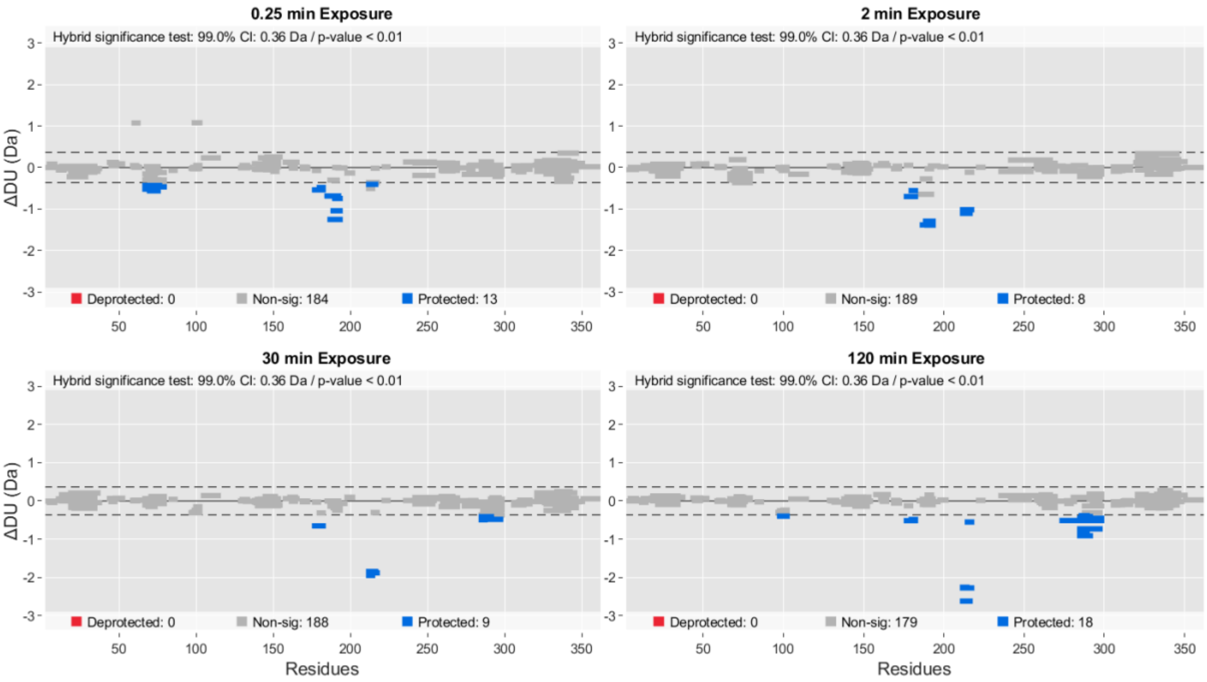

#### tβ1AR apo vs tβ1AR + Cyanopindolol

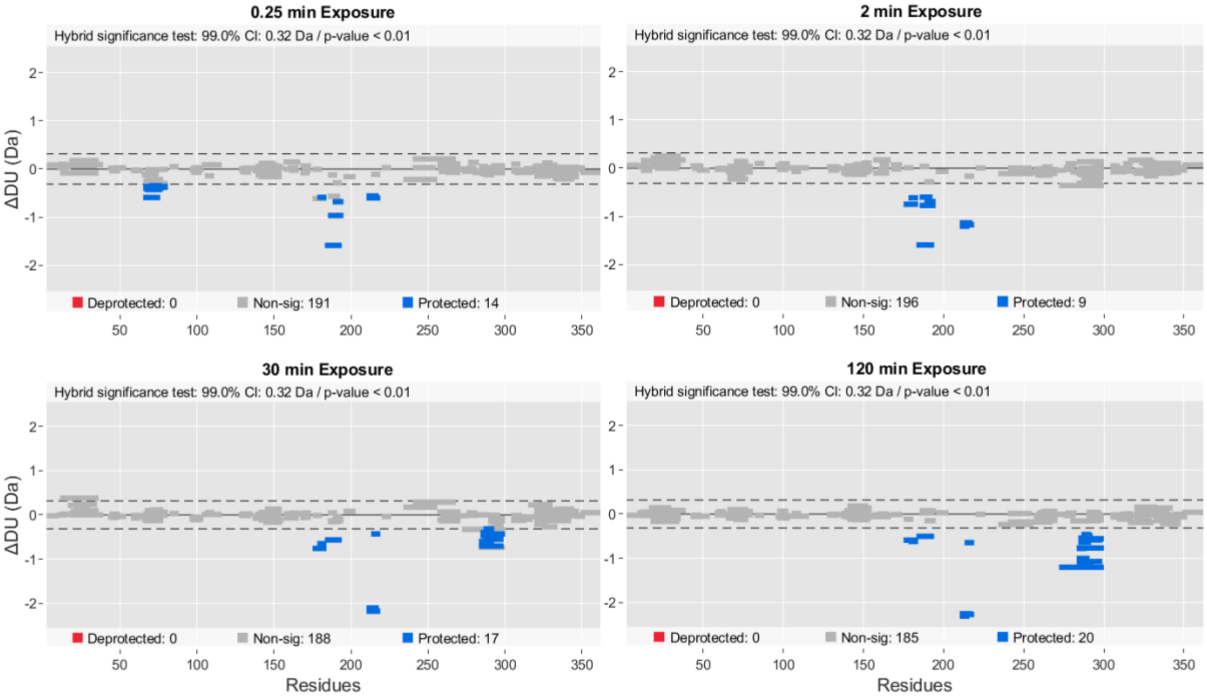


#### tβ1AR apo vs tβ1AR + Isoprenaline

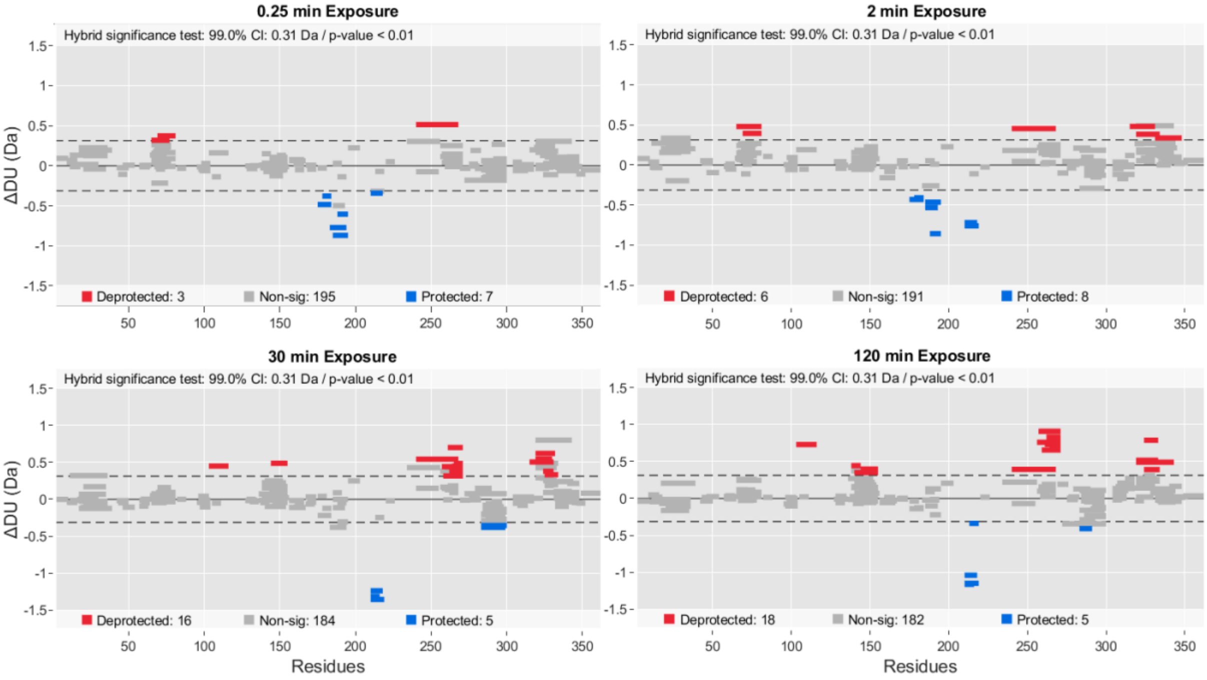

#### tβ1AR apo vs tβ1AR + Norepinephrine

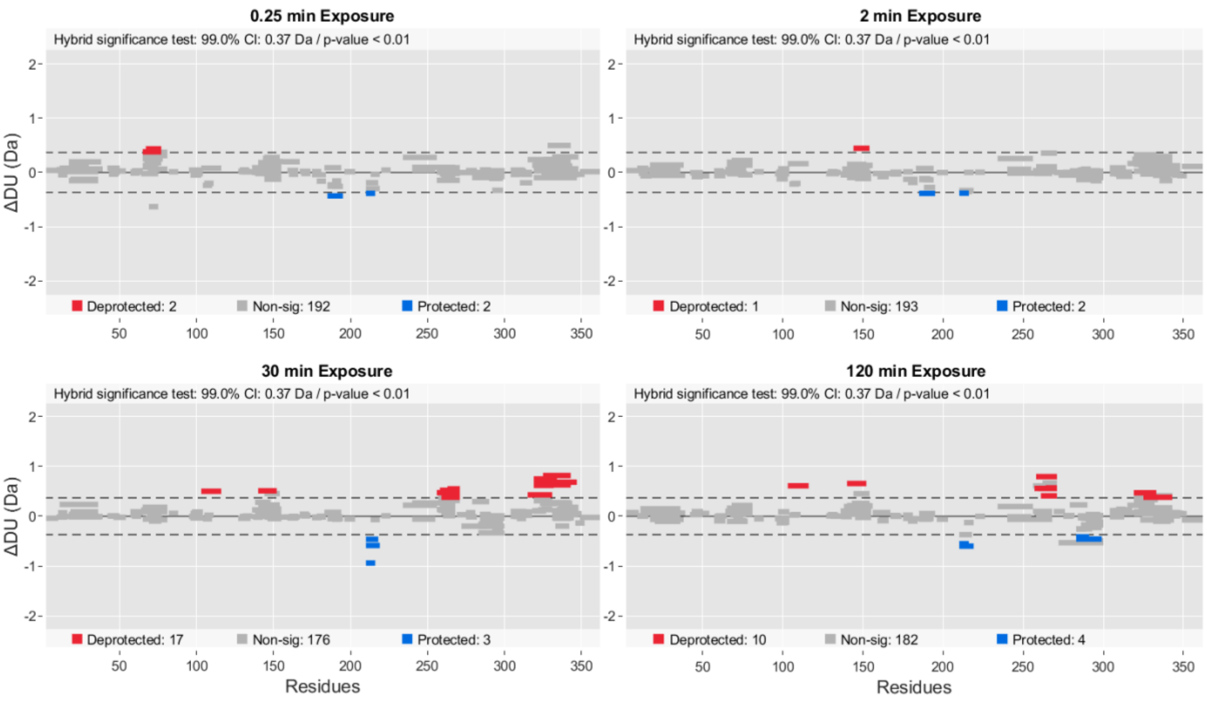


#### tβ1AR apo vs tβ1AR + Dobutamine

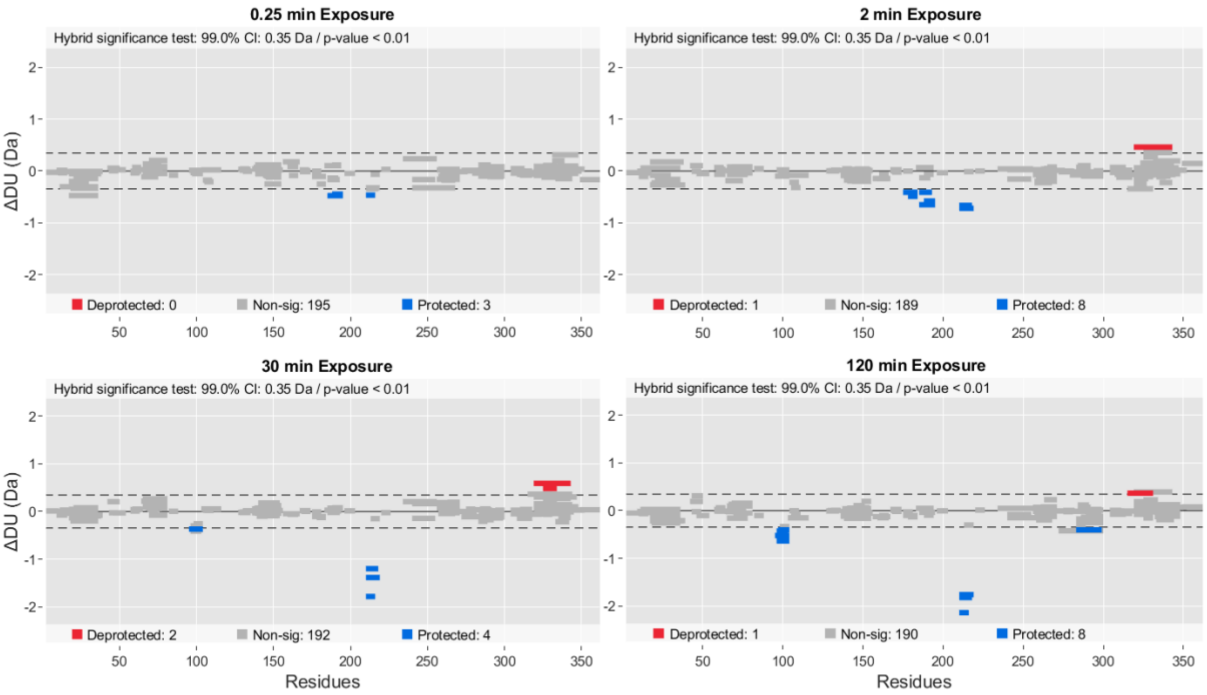

**tβ1AR apo vs tβ1AR + Salbutamol**

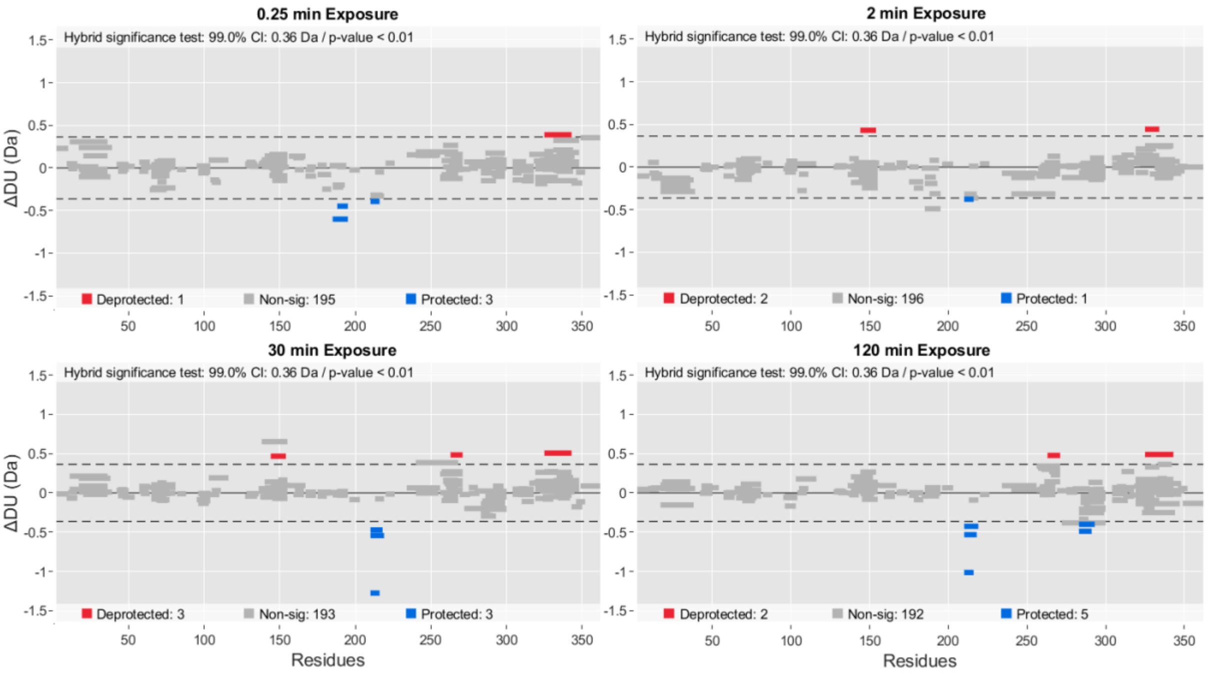


**Supplementary Figure 7. HDX Experiment 1: tβ1AR with ligands (agonist, partial agonist, antagonist).** Deuterium uptake differences presented in Woods plot, which provides breakdown of peptides ensemble for 0.25-, 2-, 30- and 120-min time point. It displays peptide length, global coverage, and deuterium uptake. 99% confidence limit was applied to the data set to identify peptides with significant deuterium uptake. Deprotected, protected and non-significantly different peptides are in red, blue, and grey respectively. Presented plots have been created in Deuteros 2.0[33].

**tβ1AR apo vs tβ1AR + Orciprenaline**

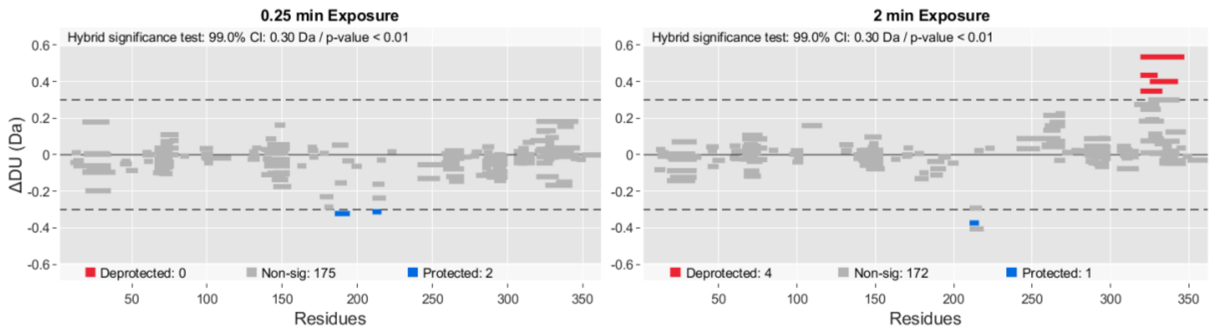


**tβ1AR apo vs tβ1AR + Isoprenaline**

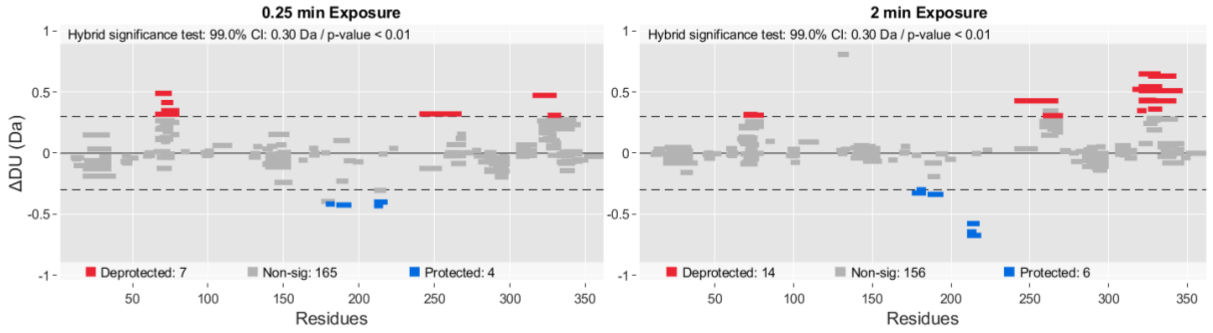


**tβ1AR apo vs tβ1AR + Colterol HCl**

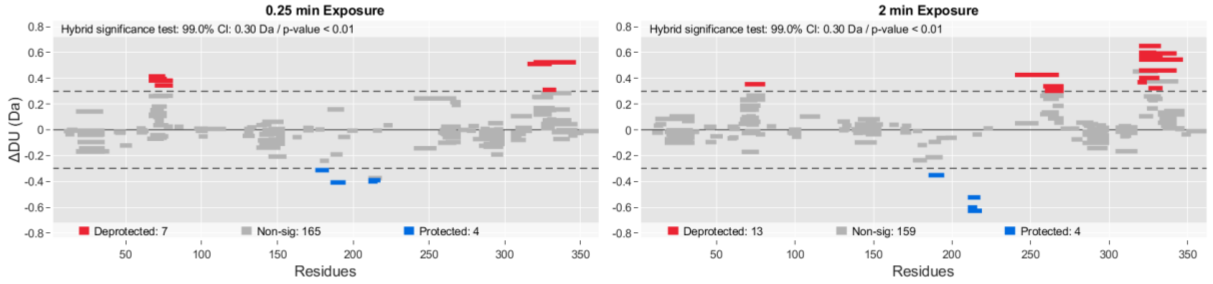


**Supplementary Figure 8. HDX Experiment 2: tβ1AR with Isoprenaline derivatives.**

Deuterium uptake differences presented in Woods plot, which provides breakdown of peptides ensemble for 0.25- and 2-min time point. It displays peptide length, global coverage, and deuterium uptake. 99% confidence limit was applied to the data set to identify peptides with significant deuterium uptake. Deprotected, protected and non-significantly different peptides are in red, blue, and grey respectively. Presented plots have been created in Deuteros 2.0[33].

#### tβ1AR_L72A vs tβ1AR_L72A + Cyanopindolol

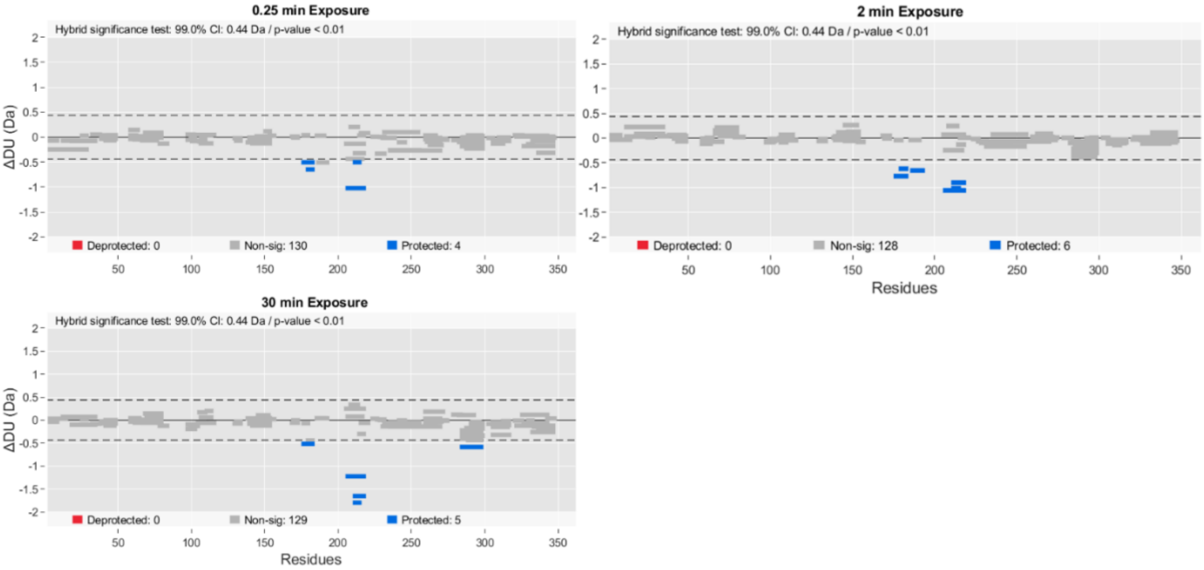


#### tβ1AR_L72A vs tβ1AR_L72A + Isoprenaline

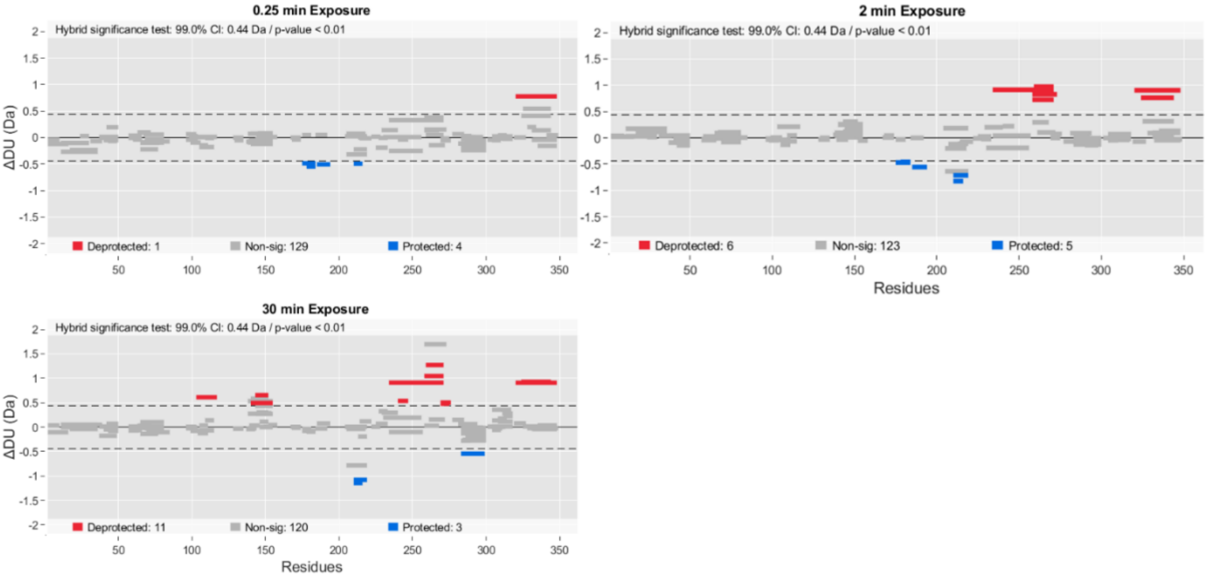


**Supplementary Figure 9. HDX Experiment 3: tβ1AR_L72A mutant with Isoprenaline and Cyanopindolol.**

Deuterium uptake differences presented in Woods plot, which provides breakdown of peptides ensemble for 0.25-, 2- and 30- min time point. It displays peptide length, global coverage, and deuterium uptake. 99% confidence limit was applied to the data set to identify peptides with significant deuterium uptake. Deprotected, protected and non-significantly different peptides are in red, blue, and grey respectively. Presented plots have been created in Deuteros 2.0[33].

#### tβ1AR apo vs tβ1AR + miniGs + Isoprenaline

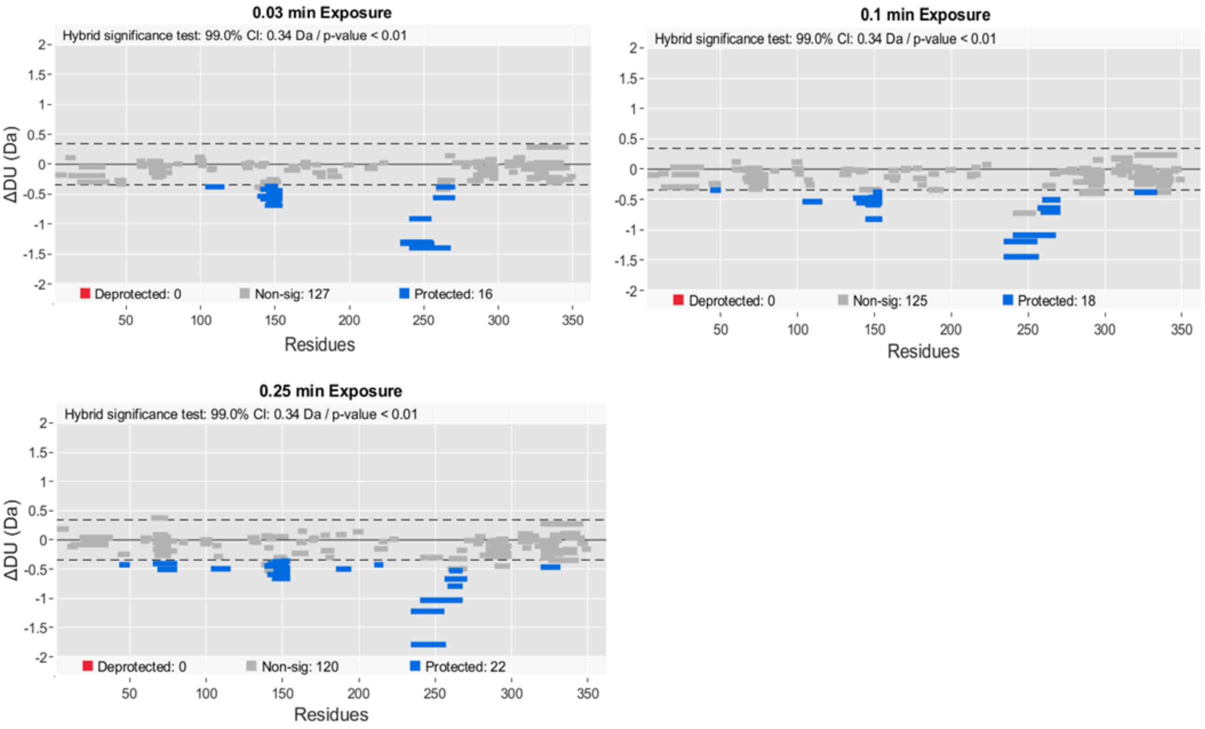


#### tβ1AR apo vs tβ1AR + miniGs + Dobutamine

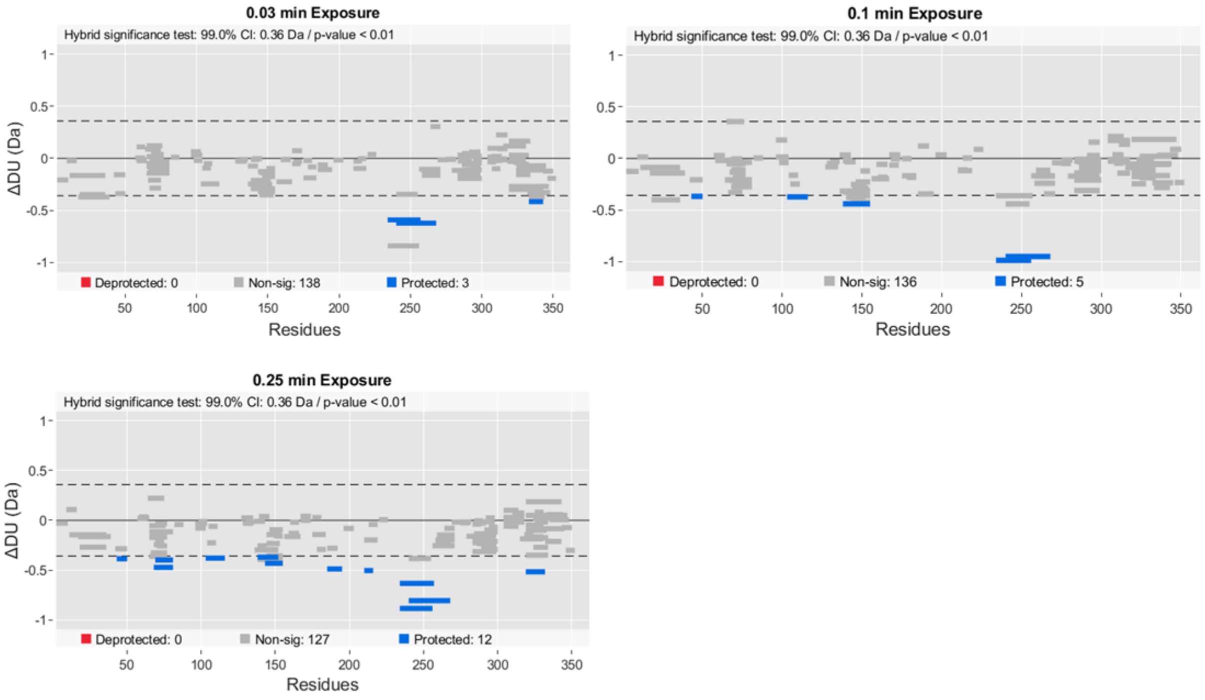

#### tβ1AR vs tβ1AR + miniGs + Cyanopindolol

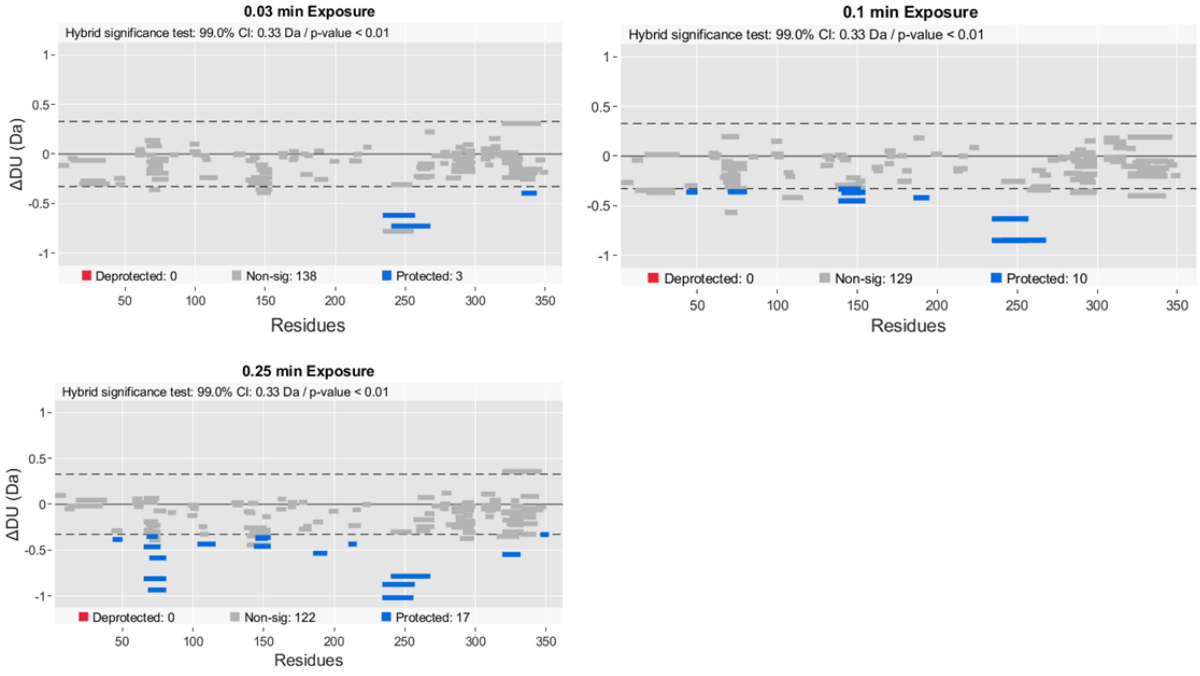


#### miniGs alone vs tβ1AR + miniGs + Isoprenaline

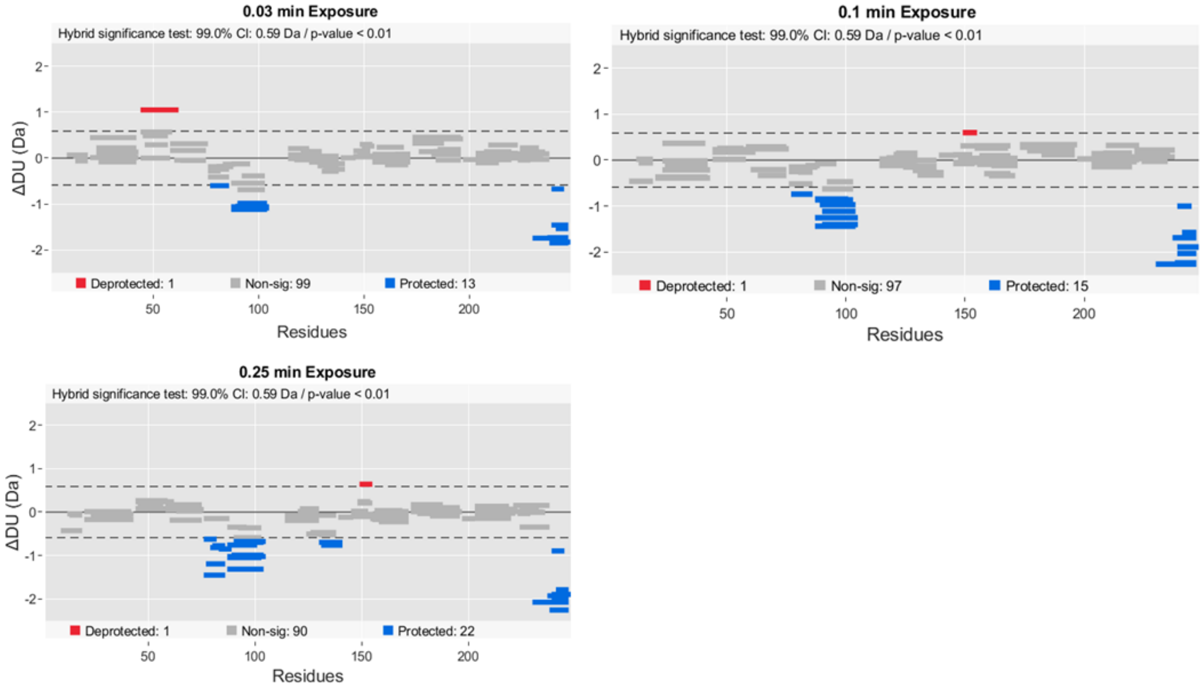

#### miniGs alone vs tβ1AR + miniGs + Dobutamine

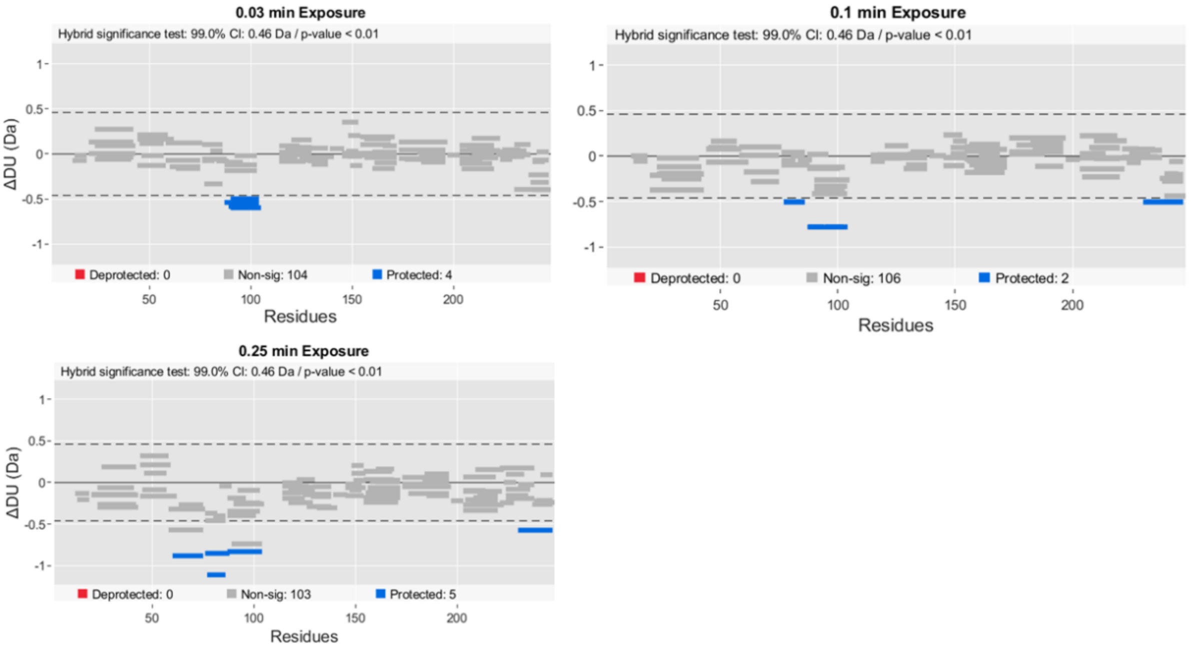


#### miniGs alone vs tβ1AR + miniGs + Cyanopindolol

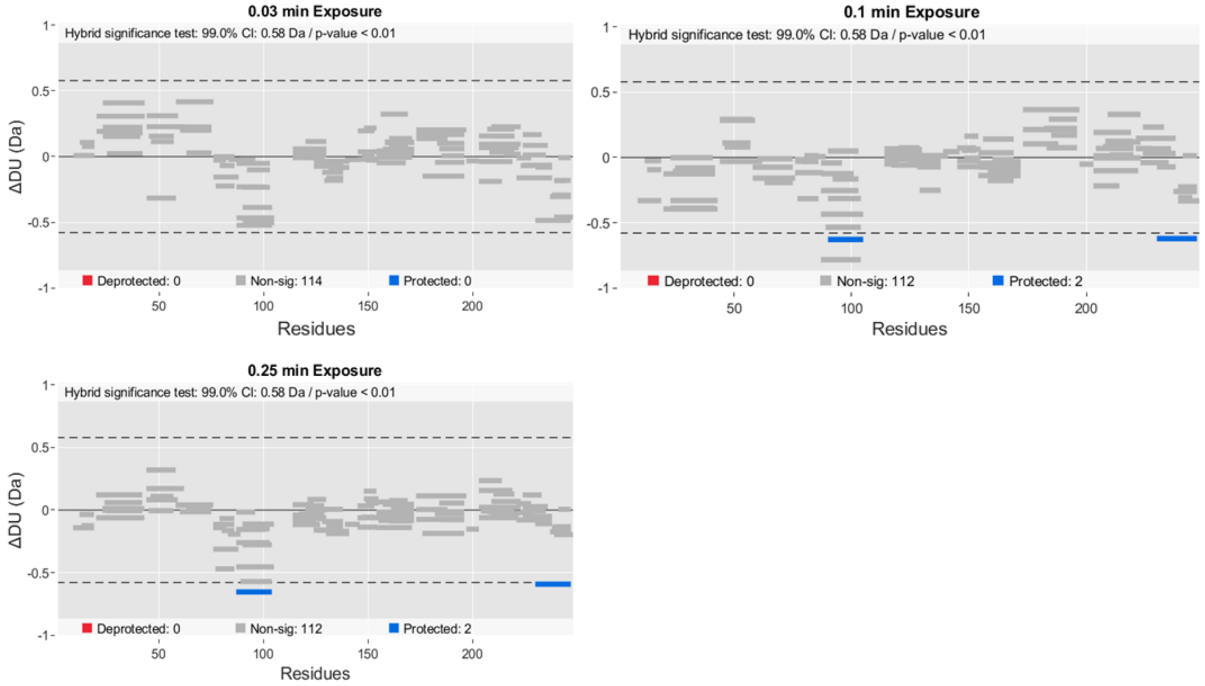


**Supplementary Figure 10. HDX Experiment 4: tβ1AR in complex with miniGs and Isoprenaline, Dobutamine and Cyanopindolol.**

Deuterium uptake differences presented in Woods plot, which provides breakdown of peptides ensemble for 0.025-, 0.1-, 0.25 min time point. It displays peptide length, global coverage, and deuterium uptake. 99% confidence limit was applied to the data set to identify peptides with significant deuterium uptake. Deprotected, protected and non-significantly different peptides are in red, blue, and grey respectively. Presented plots have been created in Deuteros 2.0[33].

**tβ1AR L72A alone vs tβ1AR L72A + miniGs + Isoprenaline**

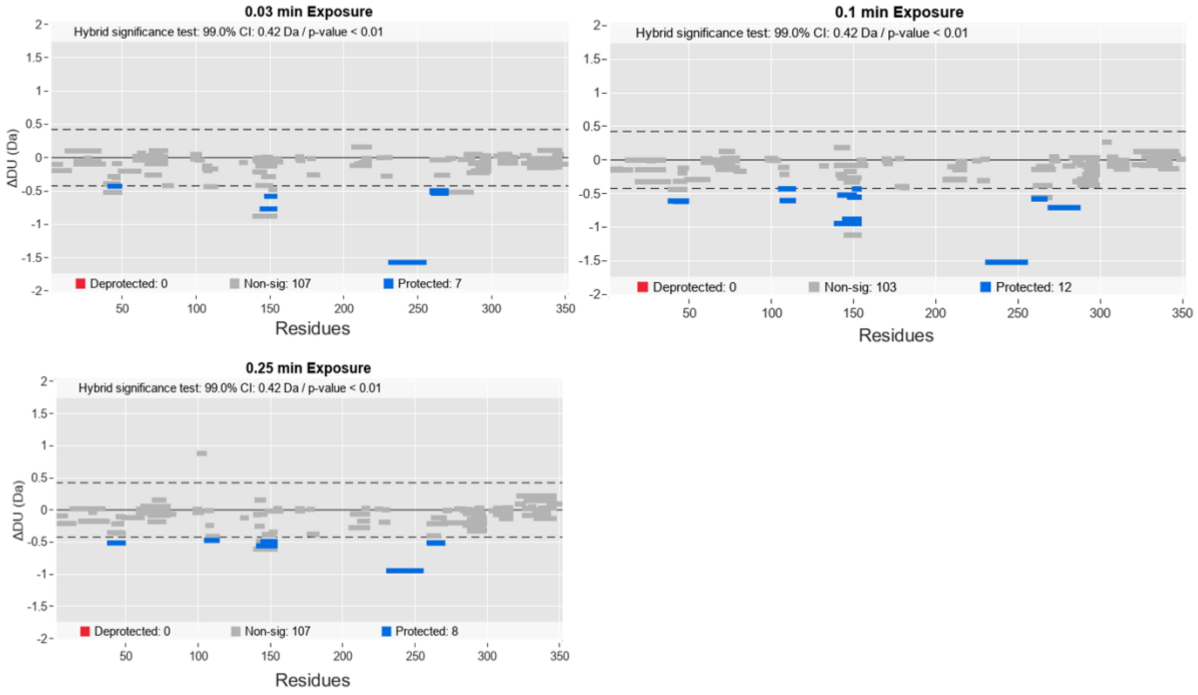


**miniGs alone vs tβ1AR L72A + miniGs + Isoprenaline**

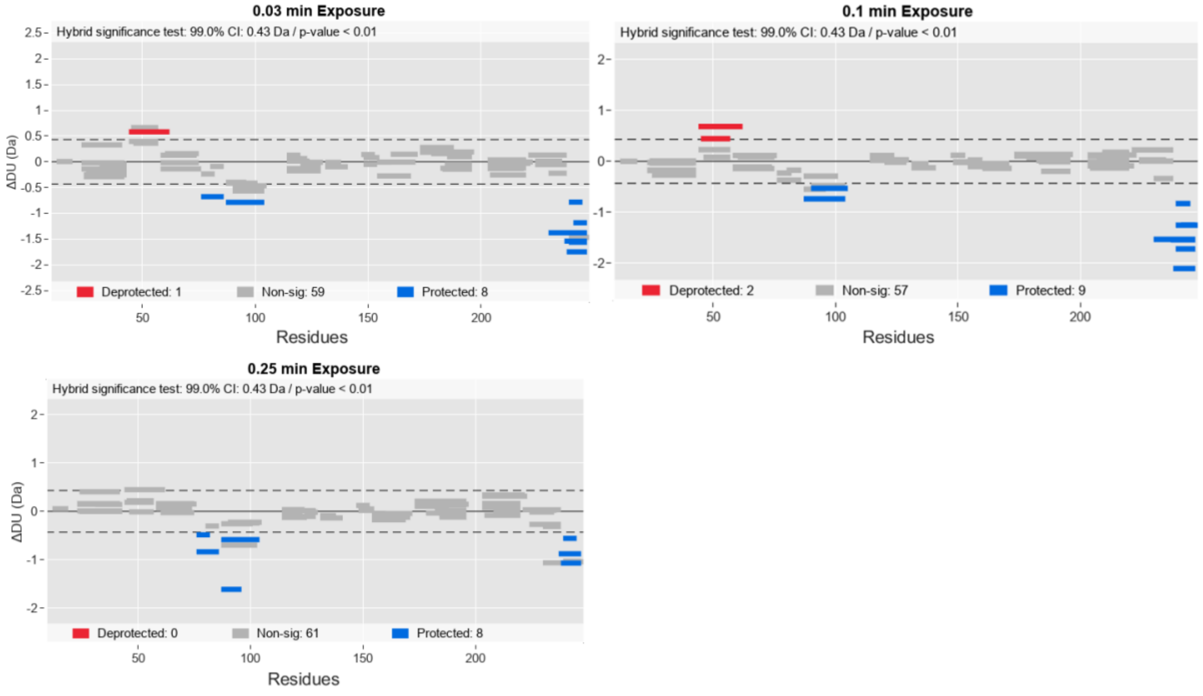


**Supplementary Figure 11. HDX Experiment 5: tβ1AR L72A mutant in complex with miniGs and Isoprenaline.**

Deuterium uptake differences presented in Woods plot, which provides breakdown of peptides ensemble for 0.025-, 0.1-, 0.25 min time point. It displays peptide length, global coverage, and deuterium uptake. 99% confidence limit was applied to the data set to identify peptides with significant deuterium uptake. Deprotected, protected and non-significantly different peptides are in red, blue, and grey respectively. Presented plots have been created in Deuteros 2.0[33].

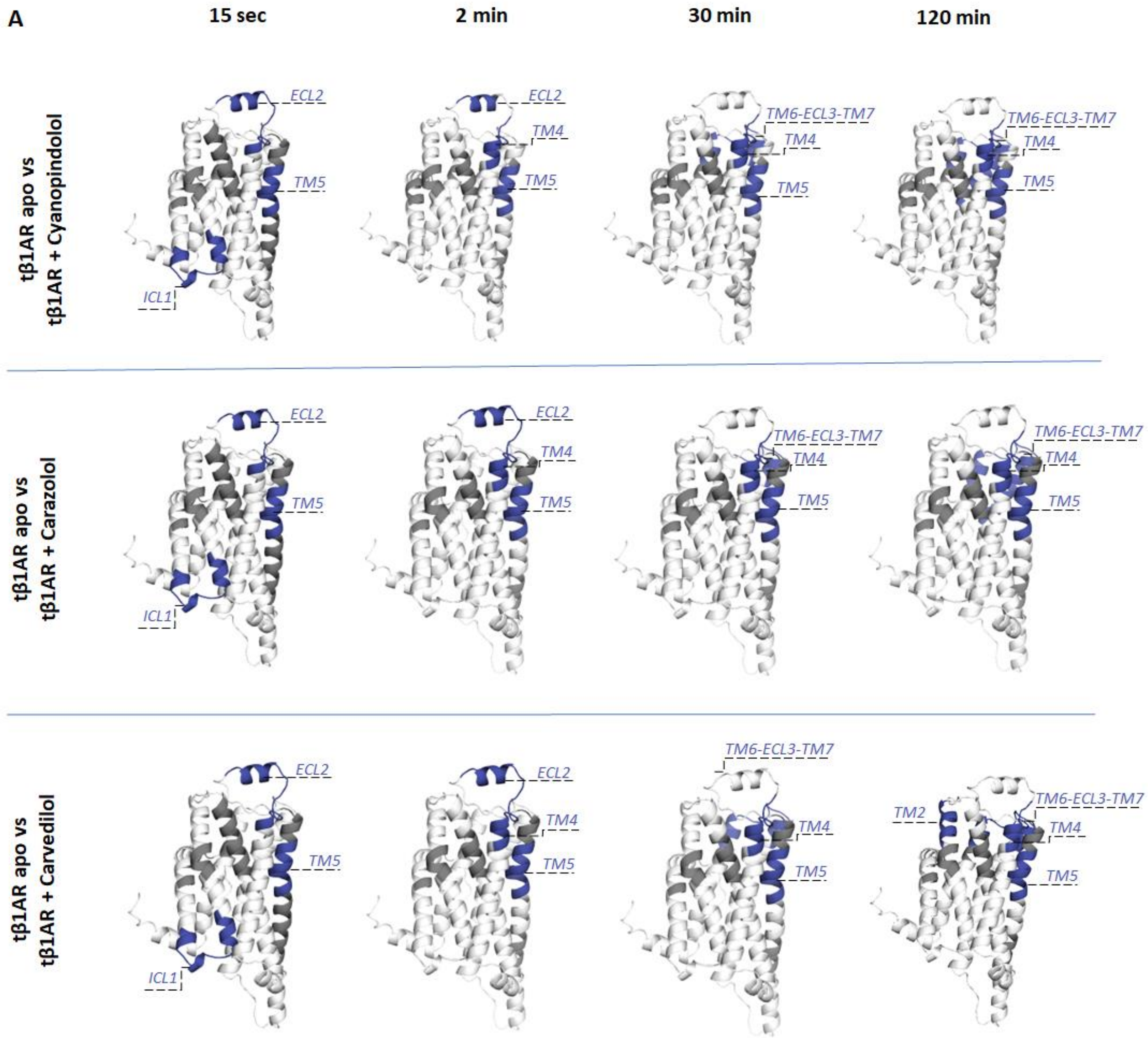
A
15 sec
2 min
30 min
120 min
tβ1AR apo vs
tβ1AR + Cyanopindolol
tβ1AR apo vs
tβ1AR + Carazolol
tβ1AR apo vs
tβ1AR + Carvedilol
ECL2
TM4
TM5
ICL1
TM6-ECL3-TM7
TM2

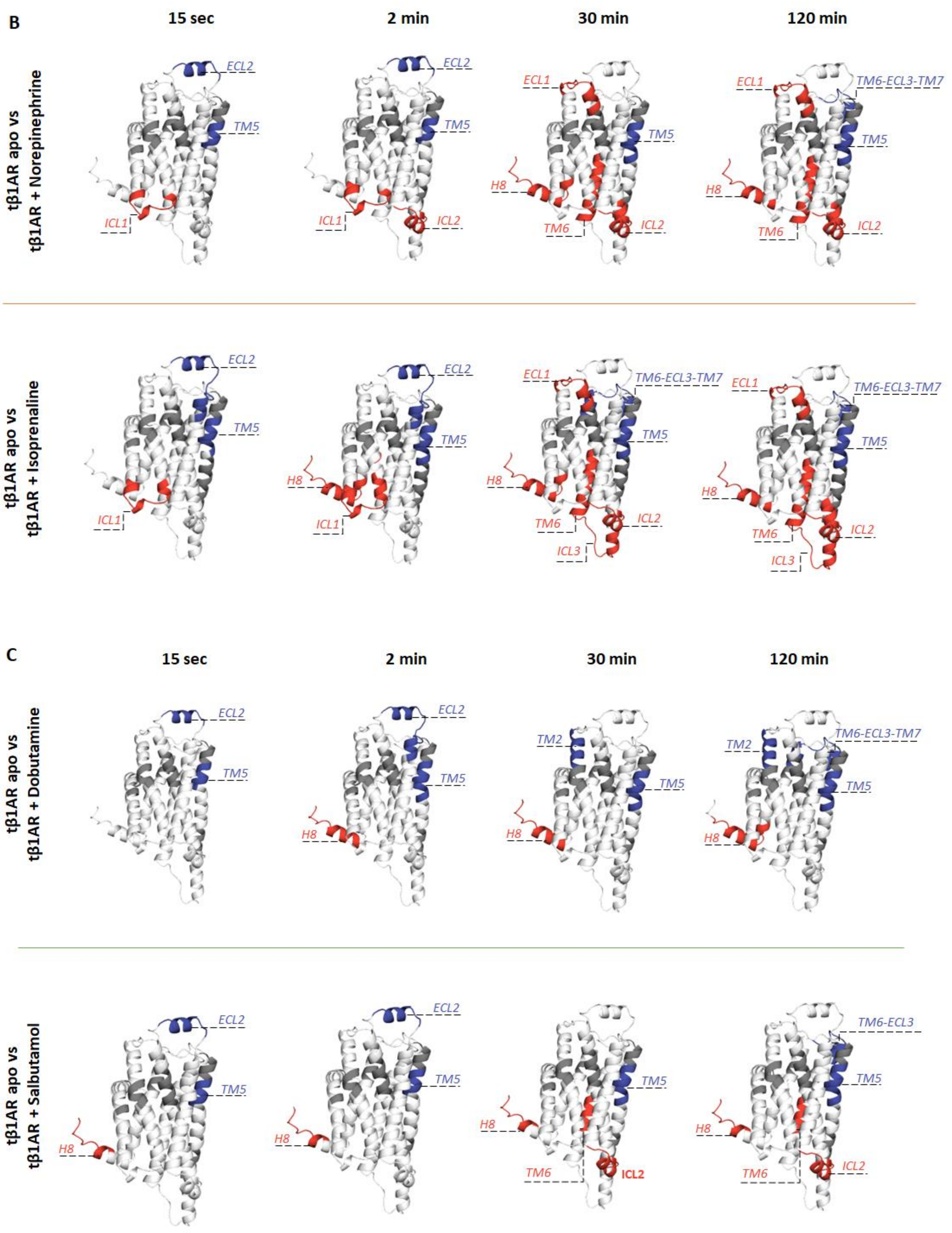


**Supplementary Figure 12. HDX Experiment 1. tβ1AR with ligands (agonist, partial agonist, antagonist).**

Graphical visualisation of the results obtained from the experiment 1. **A** Results mapped onto β1AR crystal structure (modelled by use of PDB structures; 2VT4 chain A and 6IBL chain A) for all tested antagonists. **B** Results for all tested agonists. **C** Results for all tested partial agonists. Differential deuterium uptake is plotted for each time point (15 sec, 2min, 30 min, 120 min). Red and blue indicate deprotection and protection respectively. Figures were created in PyMOL.

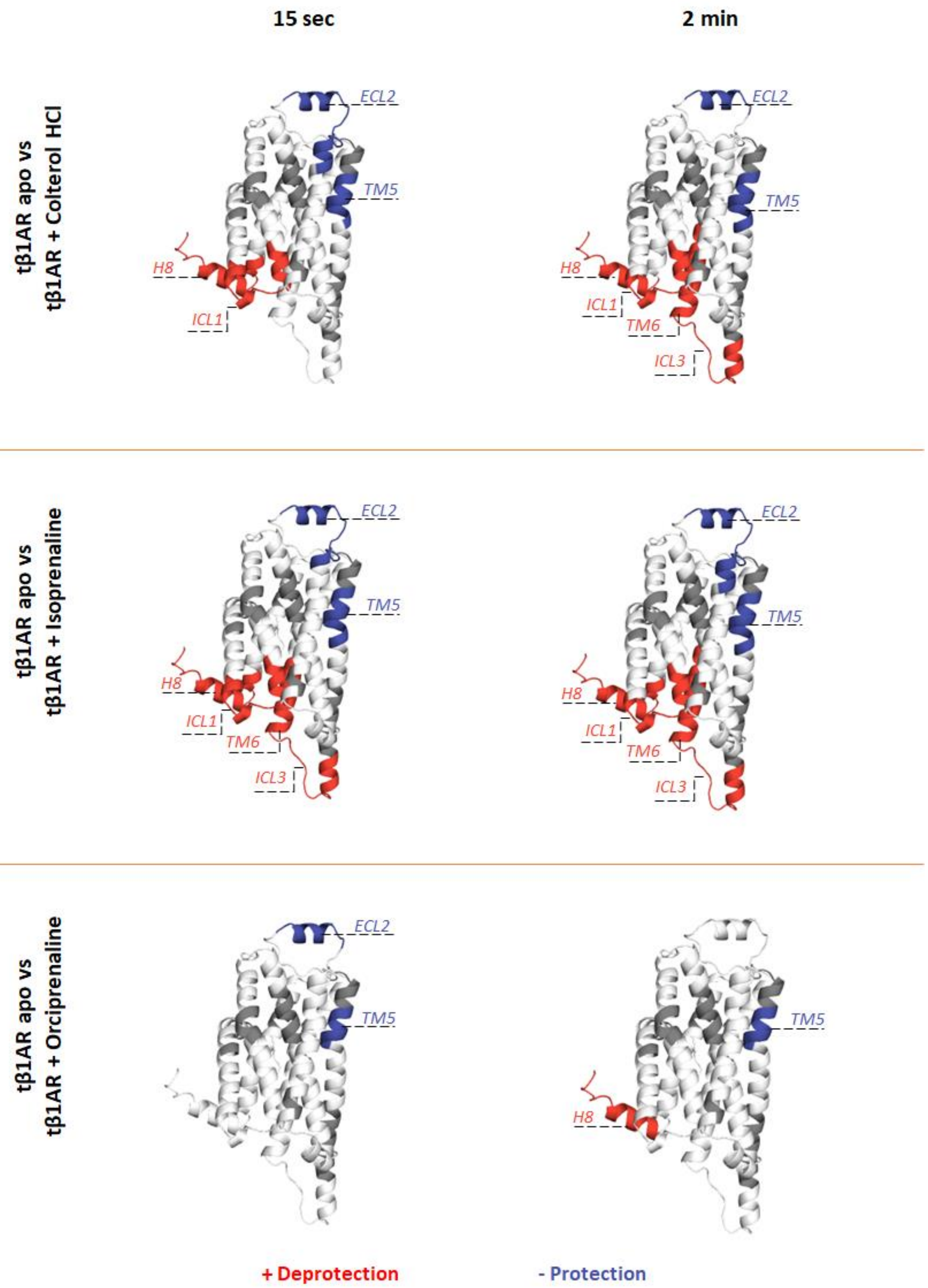


**Supplementary Figure 13. HDX Experiment 2: tβ1AR with Isoprenaline derivatives.**

Graphical visualisation of the results obtained from the experiment 2. Observed effects for tβ1AR bound to Colterol HCl, Isoprenaline and Orciprenaline are mapped onto β1AR crystal structure (modelled by use of PDB structures; 2VT4 chain A and 6IBL chain A). Differential deuterium uptake is plotted for each time point (15 sec, 2min). Red and blue indicate deprotection and protection respectively. Figures were created in PyMOL.

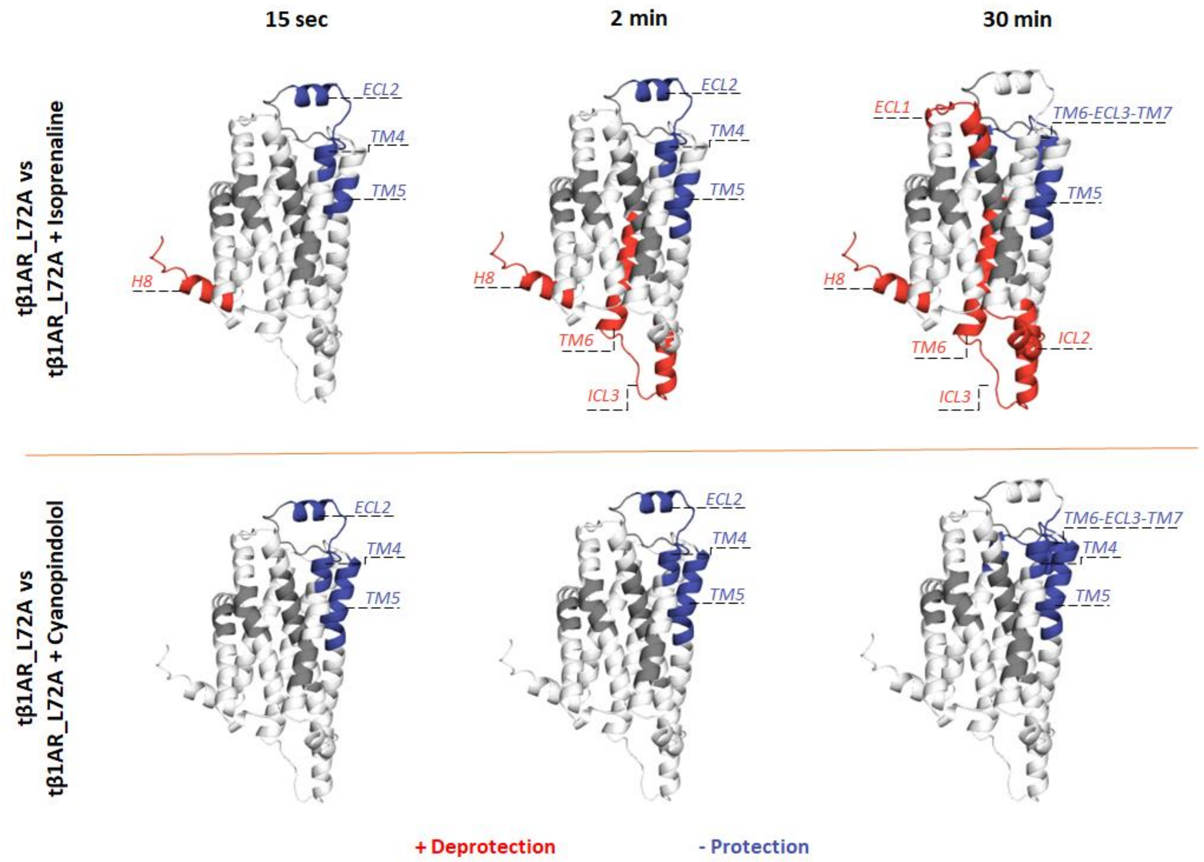


**Supplementary Figure 14. HDX Experiment 3: tβ1AR_L72A mutant with Isoprenaline and Cyanopindolol.**

Graphical visualisation of the results obtained from the experiment 3. Observed effects for tβ1AR_L72A mutant bound to agonist isoprenaline and antagonist cyanopindolol are mapped onto β1AR crystal structure (modelled by use of PDB structures; 2VT4 chain A and 6IBL chain A). Differential deuterium uptake is plotted for each time point (15 sec, 2min, 30 min). Mutation of the residue L72 located on ICL1 abolished effects previously observed for that loop. Red and blue indicate deprotection and protection respectively. Figures were created in PyMOL.

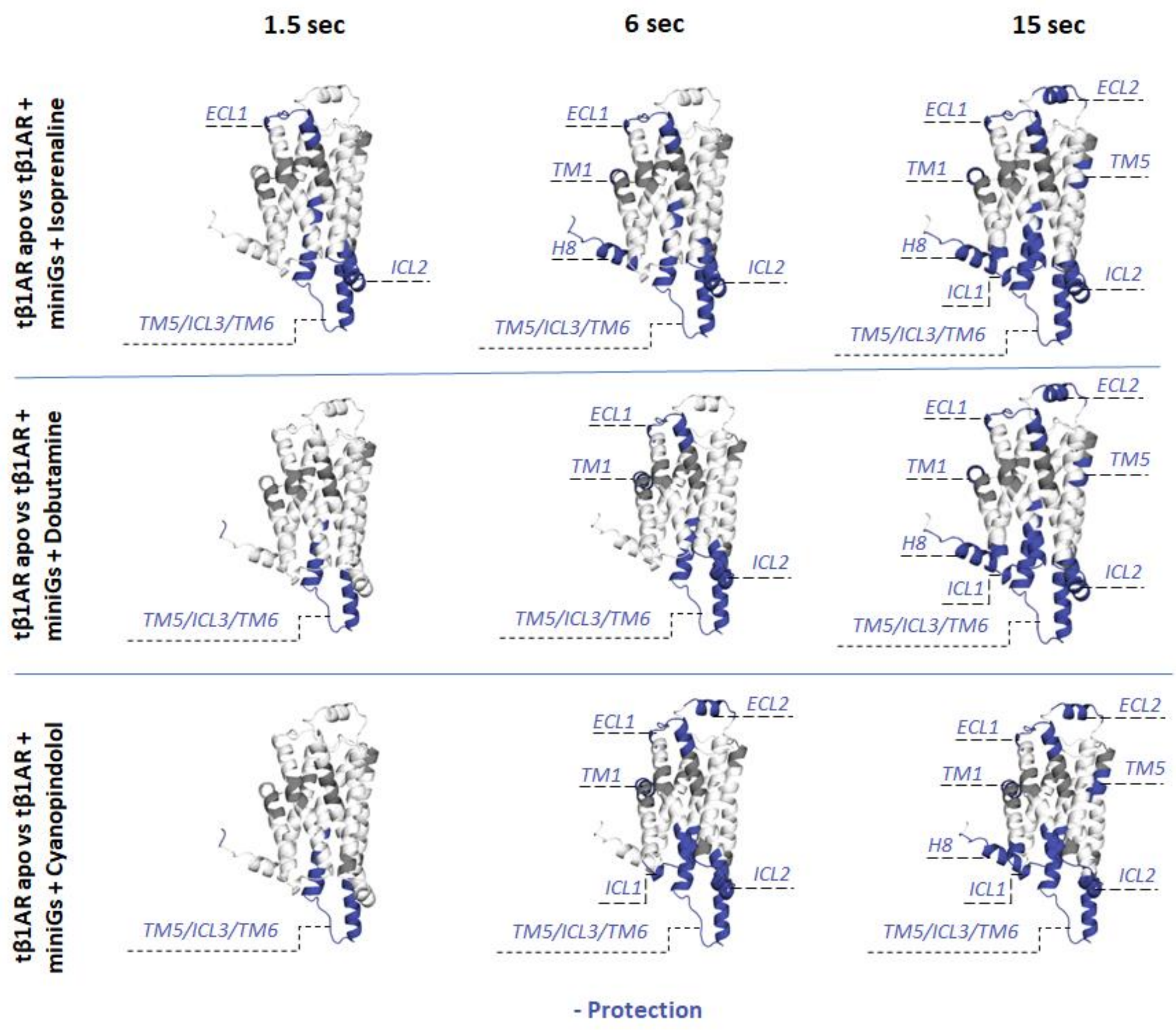
1.5 sec
6 sec
15 sec
tβ1AR apo vs tβ1AR + miniGs + Isoprenaline
tβ1AR apo vs tβ1AR + miniGs + Dobutamine
tβ1AR apo vs tβ1AR + miniGs + Cyanopindolol
ECL1
ECL2
TM1
TM5
H8
ICL1
ICL2
TM5/ICL3/TM6
- Protection

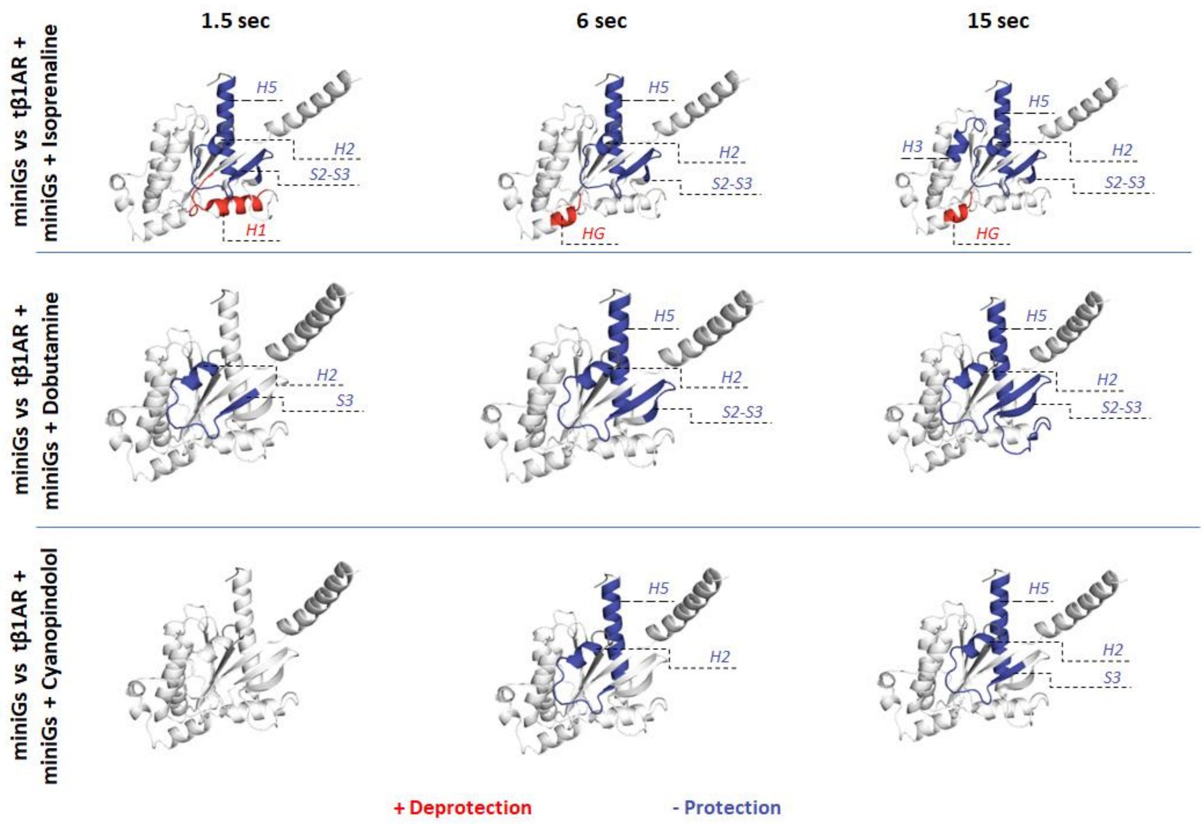


**Supplementary Figure 15. HDX Experiment 4: tβ1AR in complex with miniGs and Isoprenaline, Dobutamine and Cyanopindolol.**

Graphical visualisation of the results obtained from the experiment 4. Observed effects for tβ1AR coupled to miniGs and agonist isoprenaline, antagonist cyanopindolol and partial agonist dobutamine, are mapped onto β1AR crystal structure (modelled by use of PDB structures; 2VT4 chain A and 6IBL chain A) and miniGs crystal structure. Differential deuterium uptake is plotted for each time point (1.5 sec, 6 sec and 15 sec). Blue indicates protection and red deprotection. Figures were created in PyMOL.

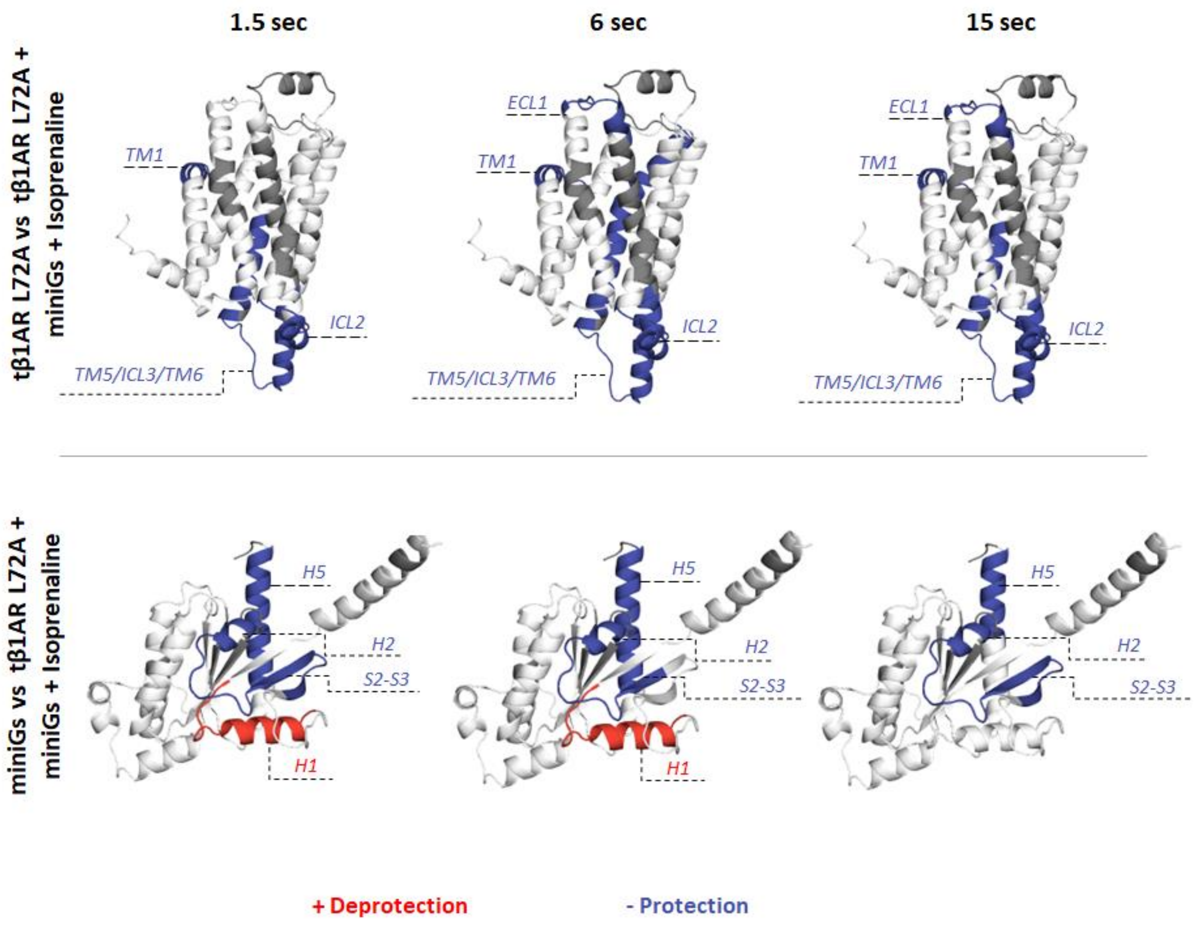


**Supplementary Figure 16. HDX Experiment 5: tβ1AR L72A mutant in complex with miniGs and Isoprenaline.**

Graphical visualisation of the results obtained from the experiment 5. Observed effects for tβ1AR L72A mutant coupled to miniGs and agonist isoprenaline are mapped onto β1AR crystal structure (modelled by use of PDB structures; 2VT4 chain A and 6IBL chain A) and miniGs crystal structure. Differential deuterium uptake is plotted for each time point (1.5 sec, 6 sec and 15 sec). Blue indicates protection and red deprotection. Figures were created in PyMOL.
